# Essential Considerations for Free Energy Calculations of RNA-Small Molecule Complexes: Lessons from Theophylline-Binding RNA Aptamer

**DOI:** 10.1101/2024.08.16.608304

**Authors:** Ali Rasouli, Frank C. Pickard, Sreyoshi Sur, Alan Grossfield, Mehtap Işık Bennett

## Abstract

Alchemical free energy calculations are widely used to predict the binding affinity of small molecule ligands to protein targets; however, the application of these methods to RNA targets has not been deeply explored. We systematically investigated how modeling decisions affect the performance of absolute binding free energy calculations for a relatively simple RNA model system: theophylline-binding RNA aptamer with theophylline and five analogs. The goal of this investigation was twofold: (1) understanding the performance levels we can expect from absolute free energy calculations for a simple RNA complex and (2) learning about practical modeling considerations that impact the success of RNA binding predictions, which may be different than the best practices established for protein targets. We learned that magnesium ion (Mg^2+^) placement is a critical decision that impacts affinity predictions. When information regarding Mg^2+^ positions is lacking, implementing RNA backbone restraints is an alternative way of stabilizing RNA structure that recapitulates prediction accuracy. Since mistakes in Mg^2+^ placement can be detrimental, omitting magnesium ions entirely and using RNA backbone restraints is attractive as a risk-mitigating approach. We found that predictions are sensitive to modeling experimental buffer conditions correctly, including salt type and ionic strength. We explored the effects of sampling in the alchemical protocol, choice of the ligand force field (GAFF2/OpenFF Sage), and water model (TIP3P/OPC) on predictions, which allowed us to give practical advice for the application of free energy methods to RNA targets. By capturing experimental buffer conditions and implementing RNA backbone restraints, we were able to compute binding affinities accurately (MAE = 2.2 kcal/mol, Pearson’s correlation coefficient = 0.9, Kendall’s tau = 0.7). We believe there is much to learn about how to apply free energy calculations for RNA targets and how to enhance their performance in prospective predictions. This study is an important first step for learning best practices and special considerations for RNA-ligand free energy calculations. Future studies will consider increasingly complicated ligands and diverse RNA systems and help the development of general protocols for therapeutically relevant RNA targets.

## 1 Introduction

In recent years, the use of RNA as a therapeutic modality has garnered significant attention and has shown promising clinical success [1–3]. RNA can serve as the active molecule designed to create the therapeutic effect or it can be the therapeutic target of a small molecule binder with the ability to inhibit or modulate [4]. These applications create a motivation for developing computer-aided design approaches for RNA binders and modeling RNA-ligand interactions.

For mRNA-based therapeutics, designing efficient formulations is crucial for the potency, stability, cellular uptake, and translation efficiency [5, 6]. mRNA lipid nanoparticle formulations have a diverse composition of excipients including lipids, polymers, and small organic molecules with diverse functions including buffering and cryoprotection [7]. We believe that exploring the interaction between formulation components and specific regions of RNA has the potential to guide the design of future formulations. RNA-targeting small molecule inhibitor design is an obvious application of RNA-small molecule binding affinity predictions, but accurately estimating the binding free energies between RNA and small molecules can also guide the design of excipients and formulation components to achieve better drug product characteristics, especially when fine-tuning their RNA affinity might be required.

Binding free energy calculations; using all-atom molecular dynamics (MD) simulations and alchemical free energy methods; have emerged as powerful tools in understanding and predicting ligand-receptor interactions [8–13]. Al-chemical methods, such as free energy perturbation (FEP) [14–16] and thermodynamic integration (TI) [17–19], are widely used in computer-aided drug discovery for targeting proteins with small molecule inhibitors [20, 21].

However, using alchemical free energy calculations to predict the RNA-binding affinity of small molecules is uncommon and relatively uncharted territory. There are only a handful of studies that report relative free energy calculations for RNA and these are limited to model systems, not clinical targets [22–24]. Some of these studies focus on predicting the effect of RNA mutations on on binding free energies, instead of predicting relative affinities of small molecule ligands with diverse chemistry [25–27]. Chen et al.’s work on guanine riboswitch and Tanida et al.’s publications on theophylline-binding RNA aptamer are the only published examples of absolute binding free energy calculations for RNA targets to our knowledge [25, 28, 29].

In this study, we focused on absolute binding free energy calculations for RNA targets, instead of relative free energy calculations. The main reason is the potential of absolute calculations to be used for comparing predicted binding affinities of structurally unrelated ligands or ligands that assume different binding modes or even different binding sites. Although relative free energy calculations have cost and accuracy advantages for ranking binding affinities of closely related ligands, this approach is not suitable for comparing dissimilar ligands with different binding modes. For the potential application of free energy calculations to the design of formulation components and excipients of RNA therapeutics, we anticipate the need to model chemically diverse ligands that may interact with RNA binding sites in different ways. With this perspective, we focused our efforts on absolute free energy calculations, despite the challenges they bring.

The application of binding free energy calculations to RNA targets is in its infancy and there is much to learn. Working with RNA poses additional challenges compared to protein targets. The unique characteristics of RNA, including its negatively charged polymer backbone, higher conformational flexibility compared to proteins, and water-exposed binding sites, introduce significant technical challenges for modeling RNA-small molecule complexes. Moreover, RNA has a limited chemical diversity of building blocks (4 nucleotides compared to 20 amino acids). This smaller repertoire can make it harder for models to distinguish specific interactions that contribute to the binding affinity. Consequently, there is a need to explore the performance and applicability of alchemical free energy methods specifically in the context of RNA targets.

In this study, we took the first steps towards understanding if the success of alchemical free energy simulations in predicting ligand-protein interactions can translate to RNA targets. Specifically, we explored the performance of alchemical free energy methods on a simple and well-studied model system: the theophylline-binding RNA aptamer with six theophylline analogs [22, 28–30]. This model system was chosen because it has the largest experimental affinity range (5.5 kcal/mol) among its six congeneric ligands compared to other options considered. The large range of binding affinities makes this system suitable for exploring the effect of simulation parameters and setup choices on the predicted results and allows distinguishing differences in performance. An experimental 3D structure exists for theophylline bound complex of theophylline binding RNA aptamer [30, 31]. Other structurally similar ligands most likely share the same binding site. Another advantage of this system was that all six ligands have neutral charge and lack rotatable bonds. By focusing on this relatively simple model system, we aimed to evaluate the feasibility of applying these methods to RNA targets and to provide insights into the challenges that need to be addressed when modeling RNA-ligand interactions with alchemical free energy methods.

To explore how a state-of-the-art tool performs in this new challenge, we selected the BFEE2 software package [32, 33] for the setup and analysis of the absolute binding free energy calculations. BFEE2 is a versatile tool that streamlines the setup and analysis of absolute free energy calculations with both alchemical and geometric routes. It is an open-source software that supports many force fields and configuration files for NAMD [34] or GROMACS [35]. The automation of setup and post-processing minimizes errors and achieves reproducibility. Default workflows in BFEE2 were designed to be robust for diverse protein systems. The authors have demonstrated successful results for predicting binding free energy for a diverse set of systems: protein-peptide complexes and protein-small molecule complexes with buried or surface binding sites, flexible or rigid ligands, neutral or charged ligands, aqueous and membrane protein targets [32, 33]. Although the BFEE2 workflow had not yet been applied to predict binding free energies for nucleic acid targets yet, we thought it was a promising start given the rigorous statistical mechanical framework designed to work with a broad range of challenging applications of protein targets.

The alchemical free energy perturbation (FEP) route was preferable over the geometric route of BFEE2 due to multiple reasons: (1) It is suitable for both buried or shallow (solvent-accessible binding sites on the target surface) binding sites, (2) the alchemical route reduces human intervention during the free-energy calculations, compared to the geometric route. On the other hand, geometric route may be advantageous for charged ligands in solvent exposed binding pockets. For us, the alchemical route was more attractive as we aimed to adopt workflows that can potentially predict free energies of tens of ligands at a time, which is only practically feasible with a fully automated workflow.

To gain a comprehensive understanding of the factors influencing computed RNA-small molecule binding free energies, we explored various conditions and parameters beyond the ones typically considered for free energy calculations for protein targets. Specifically, we investigated the impact of different buffer conditions, including salt type (NaCl vs KCl) and concentration, the presence of magnesium ions (Mg^2+^), and RNA backbone restraints. Protein-ligand free energies are generally not particularly sensitive to buffer conditions, but due to the highly negatively charged and flexible backbone of RNA, we suspected that capturing realistic ionic strength and ionic interactions may play a more significant role. We also examined the effect of Mg^2+^ ions that bind strongly to the RNA backbone, aiming to assess their influence on the stability of the RNA-small molecule complex.

We investigated whether applying backbone restraints for the RNA backbone provides any benefit. Restraints to control the ligand pose in the binding site are typical in alchemical free energy calculations [36], but t is not typically necessary or desirable to restrain the target macromolecule when it is a protein. However, RNA targets can be much more flexible than proteins given the presence of six backbone dihedral angles per nucleotide. To address the inherent conformational flexibility of RNA, we explored the application of restraints to the RNA backbone, seeking to mitigate the challenges associated with RNA’s dynamic nature on the convergence of free energy calculations. To do this correctly, we had to adjust the thermodynamic cycle for alchemical free energy calculation to capture the contributions of RNA backbone restraints. Target backbone restraints are not typically used in absolute free energy calculations for protein complexes. The BAT python tool developed by Heinzelmann and Gilson is the only example where conformational restraints for the macromolecular target were used for absolute free energy calculations [37]. In the BAT workflow, the conformational restraints are applied to the protein with optional harmonic potential restraints on backbone dihedral angles. In our protocol, we implemented an RMSD restraint on the RNA backbone heavy atoms, to keep the conformation of the RNA close to its experimental structure.

The small molecule force fields and water models were selected considering the compatibility with the RNA force field of choice for this study: Amber OL3 [38]. Since both Amber OL3 and Generalized Amber Force Field 2 (GAFF2) [39] were developed with TIP3P water model [40], this combination gave us the best chance of compatibility of force field parameters. In addition, we evaluated the performance of different water models by comparing the widely used TIP3P water model with the OPC water model [41]. This allowed us to assess the contributions of the water model to the accuracy of our binding free energy predictions. Furthermore, we compared the performance of GAFF2 and the Open Force Field (OpenFF Sage) [42, 43] in describing the small molecule’s interactions, aiming to understand the impact of force field choice on the accuracy of our calculations.

By systematically exploring these different conditions and parameters on a simple model system, our goal was to learn the unusual considerations necessary when applying binding free energy calculations specifically to RNA targets, compared to typical protocols for protein targets. Our study provides insights into the influence of various modeling decisions on the accuracy and reliability of alchemical free energy methods when applied to RNA-small molecule binding. This investigation is a crucial first step towards learning the best practices for achieving accurate RNA binding affinity predictions. With future studies gradually expanding this investigation to diverse RNA-ligand systems with increasing complexity, we can establish the best practices for applying alchemical free energy calculations to RNA. Reliable and wide use of free energy calculations for capturing RNA-small molecule interactions would be a beneficial addition to the computer-aided drug design toolbox for facilitating the rational design of small molecules targeting RNA or excipients for RNA formulations in different therapeutic applications.

## 2 Methods

### 2.1 Model setup

The structure of this theophylline–RNA aptamer complex was originally determined using Nuclear Magnetic Resonance (NMR) spectroscopy [30]; however, for this study, we used the refined NMR structure [31] (PDB ID: 1O15, Fig 1). For theophylline analogs (1-methylxanthine, 3-methylxanthine, hypoxanthine, xanthine, and caffeine) studied here, we used the same RNA structure and swapped theophylline with the other small molecules by aligning the ring structures of the small molecules using VMD [47].

**Figure 1.**
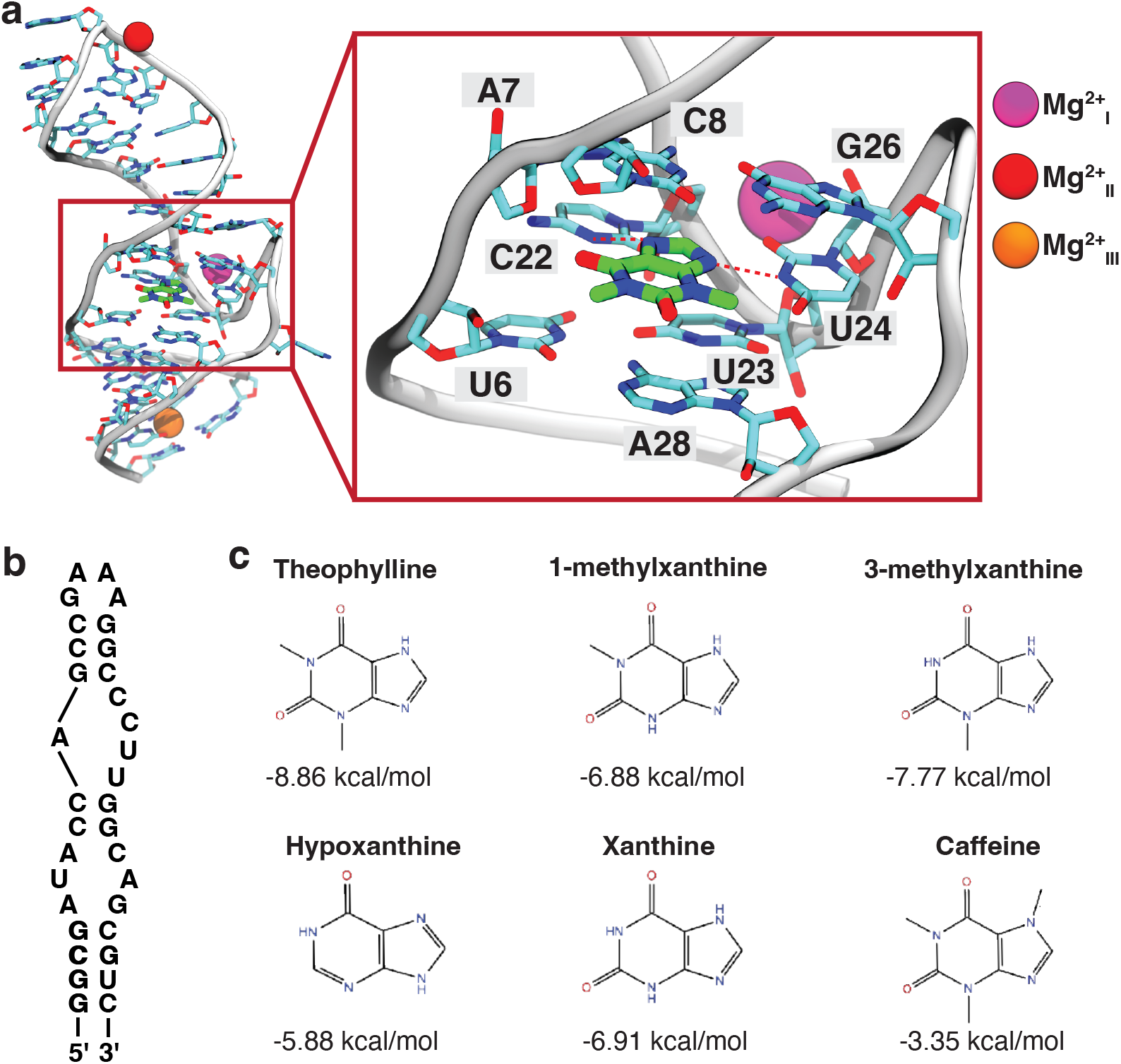
Theophylline binding RNA aptamer. **(a)** NMR structure of the theophylline binding aptamer (PDB ID: 1O15). Three Mg^2+^ ions, purple, red, and orange, were manually placed to bind the backbone of the RNA based on previous studies [22, 44, 45]. RNA side chains and theophylline are shown in licorice with carbons colored cyan and green, respectively. The zoom-in figure highlights two hydrogen bonds between theophylline and the RNA, shown as red dashed lines. **(b)** The secondary structure of the theophylline binding aptamer. **(c)** Chemical structures of theophylline and five of its analogs with experimental binding free energies [46] are shown below each compound.

To investigate the effect of structural Mg^2+^ ions on free energy calculations we set up systems with zero, two, and three Mg^2+^ ions. In systems containing two Mg^2+^ ions, the first Mg^2+^ ion 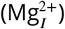 was coordinated with C22 OP1, U23 O5’, and U24 O3’ [44] and the second ion 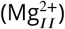 was coordinated with OP2 atoms of G14 and A15, and with OP1 atom of A16 [22]. In the three Mg^2+^ systems, a third Mg^2+^ ion 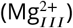 was added to coordinate with G2 O6 and U32 O4 [45]. All the Mg^2+^ ions were placed at the center of mass of the coordinating atoms. Afterward the RNA–ligand systems were solvated with a water box size of at least 68 × 68 × 68 Å^3^.

Next, we added salt to the simulation box to match the ionic strength of the binding affinity experiments, by considering the experimental buffer condition: 50 mM NaCl, 5 mM MgCl_2_, and 100 mM HEPES at pH 7.3. The ionic strength (*I*) of the experimental buffer was determined as 85 mM as follows:

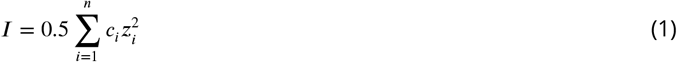

where *c*_*i*_ is the molar concentration of the *i*^th^ ion species, and *z*_*i*_ is the corresponding formal charge of the ion.

From the buffer conditions above, we calculated the following ion concentrations:

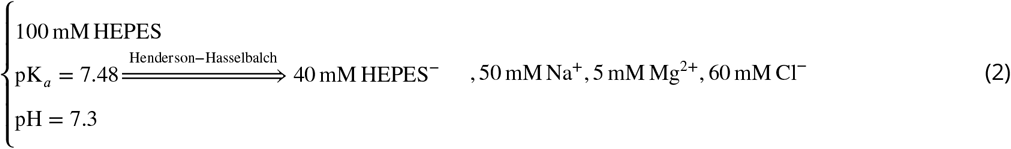

To approximate the experimental ionic strength in our simulations, we added 55 mM NaCl to the three Mg^2+^ system, yielding an ionic strength of 84 mM. Then, we used the same NaCl concentration, 55 mM, to ionize our zero and two Mg^2+^ systems as well. For comparison purposes, we prepared some systems with a higher salt concentration of 150 mM as well. Moreover, we changed the cations by ionizing some systems with KCl. In the ionization process, based on the volume of the box, we first added enough co-ion/counterion pairs to achieve the desired solution salt concentration. Next, we added more counter-ions to neutralize our simulation box. We also prepared a system in which we only neutralized the system using Na^+^ counter-ions. Table 1 lists all the conditions explored in this study. For each set of conditions, we repeated simulations three times. Each replicate simulation is independently built and simulated.

**Table 1.** Summary of the different methods explored in this study and their overall performance. Methods 1-19 indicate alchemical free energy calculations with variations in modeling conditions and parameters. Methods 20 and 21 are MM-GBSA calculations.

| Method ID | Salt Condition | # of Mg | Water Model | Ligand Force Field | # windows | RNA Backbone Restraints | MAE | RMSE | Pearson's r | Kendall's Tau | Spearman's Rho |
| --- | --- | --- | --- | --- | --- | --- | --- | --- | --- | --- | --- |
| 1 | 55 NaCl | 0 | TIP3P | GAFF2 | 40 win, 1 ns/win | No | 1.4 | 2.0 | 0.8 | 0.5 | 0.7 |
| 2 | 55 NaCl | 2 | TIP3P | GAFF2 | 40 win, 1 ns/win | No | 2.2 | 2.4 | 0.9 | 0.7 | 0.8 |
| 3 | 55 NaCl | 3 | TIP3P | GAFF2 | 40 win, 1 ns/win | No | 3.0 | 3.1 | 0.6 | 0.7 | 0.9 |
| 4 | 55 NaCl | 0 | TIP3P | GAFF2 | 40 win, 1 ns/win | Yes | 2.2 | 2.5 | 0.6 | 0.7 | 0.9 |
| 5 | 55 NaCl | 2 | TIP3P | GAFF2 | 40 win, 1 ns/win | Yes | 2.6 | 2.9 | 0.7 | 0.7 | 0.9 |
| 6 | 55 NaCl | 3 | TIP3P | GAFF2 | 40 win, 1 ns/win | Yes | 2.7 | 3.2 | 0.4 | 0.3 | 0.4 |
| 7 | 55 NaCl | 2 | OPC | GAFF2 | 40 win, 1 ns/win | No | 2.3 | 2.8 | 0.3 | 0.3 | 0.5 |
| 8 | 55 NaCl | 2 | TIP3P | OpenFF | 40 win, 1 ns/win | No | 2.4 | 2.8 | 0.5 | 0.5 | 0.7 |
| 9 | 55 NaCl | 3 | TIP3P | OpenFF | 40 win, 1 ns/win | No | 2.4 | 2.7 | 0.7 | 0.5 | 0.6 |
| 10 | 55 KCl | 2 | TIP3P | GAFF2 | 40 win, 1 ns/win | No | 1.9 | 2.5 | 0.3 | 0.1 | 0.1 |
| 11 | 55 KCl | 3 | TIP3P | GAFF2 | 40 win, 1 ns/win | No | 2.4 | 2.8 | 0.5 | 0.5 | 0.6 |
| 12 | 150 KCl | 3 | TIP3P | GAFF2 | 40 win, 1 ns/win | No | 1.7 | 2.5 | 0.2 | 0.5 | 0.6 |
| 13 | Neutralized | 3 | TIP3P | GAFF2 | 40 win, 1 ns/win | No | 2.5 | 2.7 | 0.8 | 0.5 | 0.6 |
| 14 | 55 NaCl | 2 | TIP3P | GAFF2 | 80 win, 1 ns/win | No | 2.8 | 2.9 | 0.9 | 0.7 | 0.9 |
| 15 | 55 NaCl | 2 | TIP3P | GAFF2 | 40 win, 2 ns/win | No | 2.8 | 3.0 | 0.9 | 0.6 | 0.8 |
| 16 | 55 NaCl | 2 | TIP3P | GAFF2 | 80 win, 1 ns/win | Yes | 2.2 | 2.4 | 0.7 | 0.9 | 0.9 |
| 17 | 55 NaCl | 2 | TIP3P | GAFF2 | 40 win, 2 ns/win | Yes | 2.4 | 2.6 | 0.6 | 0.6 | 0.7 |
| 18 | 55 NaCl | 0 | TIP3P | GAFF2 | 80 win, 1 ns/win | Yes | 1.8 | 2.0 | 0.8 | 0.7 | 0.9 |
| 19 | 55 NaCl | 0 | TIP3P | GAFF2 | 40 win, 2 ns/win | Yes | 2.3 | 2.6 | 0.5 | 0.7 | 0.9 |
| 20 | 55 NaCl | 2 | TIP3P | GAFF2 | MM-GBSA | No | 8.3 | 8.5 | 0.3 | 0.5 | 0.6 |
| 21 | 55 NaCl | 0 | TIP3P | GAFF2 | MM-GBSA | No | 7.8 | 8.6 | 0.1 | 0.5 | 0.4 |

Please refer to SI Section 8.1.1 for step-by-step protocol for model setup.

### 2.2 Initial equilibration simulation conditions

All systems were initially equilibrated with the following protocol unless stated otherwise: (1) 5,000 steps of energy minimization, followed by 4 ns of restrained equilibration by applying harmonic positional restraints (k = 5 kcal·mol^−1^·Å^−2^) to all RNA and ligand heavy atoms as well as the Mg^2+^ ions (if present in the system); (2) Gradual removal of all positional restraints (stepwise decreasing the restraint force constant from 5 kcal·mol^−1^·Å^−2^ to 0 kcal·mol^−1^·Å^−2^) duringa 5 ns simulation; (3) 100 ns of unrestrained equilibrium simulation. Steps 1 to 3 were all performed using NAMD3 [34, 48] in the isothermal-isobaric (NPT) ensemble at 298 K and 1 atm.

Please refer to SI Section 8.1.2 for the detailed protocol for pre-BFEE2 equilibrium simulations.

### 2.3 Simulation parameters

Unless stated otherwise, all simulations were run using NAMD3 and all free-energy calculations were performed using the following protocol: RNA was modeled using all-atom Amber OL3 (ff99bsc0χOL3) force field [38], while ligands were represented by the second generation General Amber Force Field (GAFF2), as implemented in Antechamber [39] or OpenFF 2.0.0, Sage [42, 43]. TIP3P or OPC water model were used [40, 41]. Monovalent and divalent ions were modeled with Li and Merz (12-6) ion parameters for the TIP3P water model (12-6 normal usage set) [49, 50].

A 9 Å cutoff was used for all short-range non-bonded interactions with switching starting at 8 Å for Lennard-Jones interactions. Particle mesh Ewald (PME) was used to calculate the long-range electrostatic interactions using fourth-order b-spline interpolation and 1 Å grid spacing [51]. The SHAKE algorithm was used to constrain the length of all the hydrogen-containing covalent bonds [52]. A Langevin thermostat maintained the temperature at 298 K, using a damping coefficient of 1 ps^−1^. The Nosé–Hoover Langevin piston-barostat maintained 1 atm pressure using a piston period and decay of 200 and 100 fs, respectively [53, 54]. We used a 2 fs simulation time step in all simulations.

All short-range, non-bonded forces (Lennard-Jones) were recalculated every time step, while long-range electrostatics were updated every other time step, using the r-RESPA multiple time-step algorithm [55]. Free energy estimates for each replicate were extracted using the post-treatment procedure of BFEE2 protocol with FEP estimator [56].

### 2.4 Free energy simulations

Alchemical free energy methods calculate the binding free energy of a ligand to an RNA target without extensively sampling multiple binding and unbinding events, which are beyond the reach of all-atom MD simulations. Instead, they employ a thermodynamic cycle (Fig. 2b and Fig. 2c) to connect the ligand-bound and unbound states, by taking the system through some unphysical states by removing the non-bonded interactions of the ligand with its environment. To maintain the conformation of the ligand similar to its native bound state during this unphysical transformation, it is necessary to apply restraints to the ligand, as described in section 2.4. These restraints contribute to the final binding free energy results, and thus, we have two extra steps in which we calculate the contribution of these restraints in the unbound and bound form, respectively.

**Figure 2.**
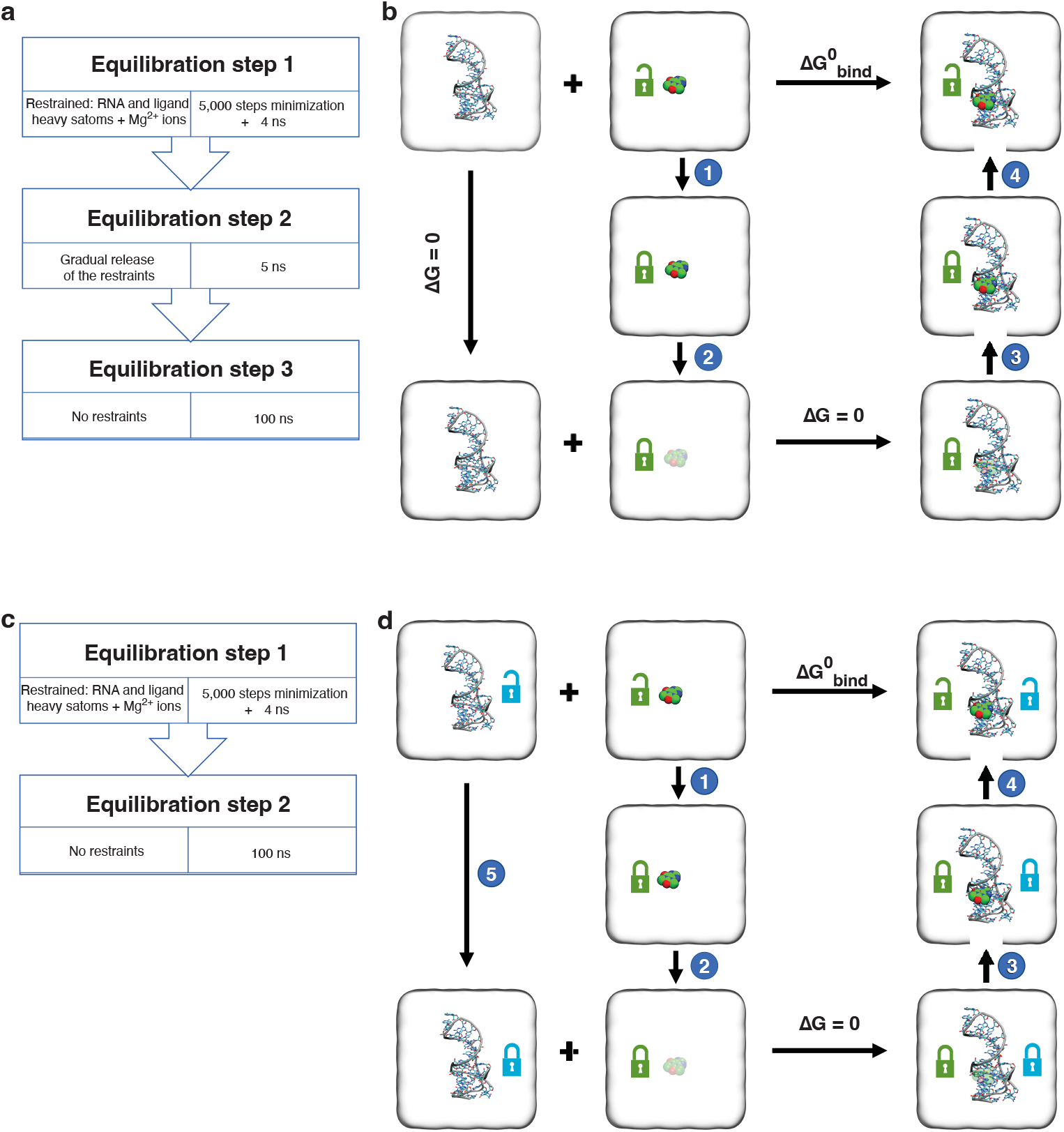
Simulation steps and thermodynamic cycle for absolute free energy calculations. (a) Step-wise pre-BFEE2 equilibration of the RNA and ligands. (b) Thermodynamic cycle based on the BFEE2 workflow [32, 56]. Open green lock represents unrestrained ligand, while closed green lock indicates application of conformational, orientational, and positional restraints on the ligand with respect to the RNA. In steps 1 and 4 free energy contributions of maintaining the ligand using restraint potentials are calculated in unbound (LlG(bulk,rest)) and bound (LlG(complex,rest)) states, respectively. Steps 2 and 3 represent the alchemical freeenergy change for reversibly decoupling the ligand from its environment in unbound (LlG(bulk,alch)) and bound (LlG(complex,alch)) states, respectively. (c) and (d) similar to **(a)** and **(b)** but for systems in which RNA backbone is restrained based on RMSD. Open and closed blue locks represent unrestrained and restrained RNA backbone, respectively. The thermodynamic cycle for free energy calculations with the RNA backbone RMSD restraints requires an extra step (step 5) to calculate the contributions of the backbone restraints. This can be calculated by simulating the RNA-only system and gradually turning constraints on and off, similar to how restraint contributions are calculated in step 4.

Alchemical free energy calculations were set up using the last frame from the third equilibration step of section 2.2 after centering the RNA and ligand in the simulation box. Using the alchemical route of BFEE2 [32, 56], simulations were set up according to the thermodynamic cycle shown in Fig. 2b. After input generation using BFEE2 GUI, a correction to ligand-only system files was necessary to take care of the extra counterions left after the removal of the RNA target with multiple negative charge. Using Ambertools we re-ionized the ligand-only systems to ensure neutrality in the absence of the RNA aptamer. The decoupling of the ligand from its environment (the RNA binding site or the bulk solution, corresponding to steps 2 and 3 of the thermodynamic cycle Fig. 2b) was calculated using free-energy perturbation (FEP) method, while its position, orientation, and conformation were restrained to its native state. To account for the energetic cost of the enforced restraints, a separate thermodynamic integration (TI) simulation was performed, in both bound and unbound systems, in which the force constants of the restraints were gradually turned off to zero (steps 1 and 4 of the thermodynamic cycle Fig. 2b). Simulations of steps 1-4 are done in a bidirectional manner to improve the reliability of the free-energy estimates [56].

Unless stated otherwise, the following parameters were used for all free energy calculations. In the bound state (RNA-ligand systems, corresponding to steps 3 and 4 of the thermodynamic cycle Fig. 2b), 40 *λ* windows were used. In the unbound state (ligand systems, corresponding to steps 1 and 2 of the thermodynamic cycle Fig. 2b), 30 *λ* windows were used. Each *λ* window was simulated for 1 ns and the first 200 ps was discarded as equilibration and not used for taking samples for free energy calculations.

In all RNA-ligand simulations, seven collective variables (CVs) were used to restrain the ligand with respect to RNA as described below [56]: **(1)** A root-mean-square deviation (RMSD) CV that keeps the ligand’s conformation restrained to its bound state native conformation (k = 10 kcal·mol^−1^·Å^−2^) **(2-3)** Standard polar and azimuthal angles (*θ, ϕ*), defined from the unit distance vector d_*unit*_ = (x,y,z) between the center of mass of the ligand and RNA (k = 0.1 kcal·mol^−1^·Å^−2^): *θ* = cos^−1^(z) and *ϕ* = atan2(y,x). These two angles specify the position of the ligand with respect to the RNA; **(4-6)** Three Euler angles defined using quaternions (q_0_, q_1_, q_2_, q_3_) that describe the best-fit rotation of the ligand with respect to its bound state (k = 0.1 kcal·mol^−1^·Å^−2^), describing orientation of the ligand; **(7)** Radial distance separating the center of mass of the ligand and that of the RNA (k = 10 kcal·mol^−1^·Å^−2^)

Please refer to SI Sections 8.1.3 and 8.5.4 for detailed BFEE2 protocol for running and analyzing alchemical free energy calculations.

### 2.5 Simulation set up with OpenFF

In one of our systems, we used OpenFF 2.0.0 (Sage) [42, 43], to describe the ligand’s interactions. To build the systems, we used the steps described in the OpenFF tutorial (see the protocol SI section 8.3). In short, we used the ParmEd python package [57] to start from a system with GAFF2 parameters for the ligand and replaced the ligand’s force field with OpenFF Sage. The RNA in this system was still described using the Amber OL3 (ff99bsc0χOL3). The TIP3P water model was used for this system, with 55 mM of NaCl and either two or three Mg^2+^ as described in section 2.1.

### 2.6 Rejection protocol for replicate quality control

In our protocol, the final estimates of the binding free energy for each ligand were reported as the mean and standard deviation of three independent replicate simulations. We adopted a replicate rejection protocol to filter out replicates with obvious quality problems. We only used replicates that passed this initial filter for calculating the final estimates of binding free energies. Low-quality replicates were detected focusing only on serious red flags obtained from simulation data: by checking for hysteresis error and free energy contribution of ligand restraints in the complex arm. The replicate rejection was performed in a principled way based on computed indicators, without making comparisons to the experimental data. Making replicate rejection decisions with an *a priori* manner was important to us for the suitability of the protocol for prospective predictions in the absence of experimental data.

To ensure all the simulated replicas are healthy we defined the following criteria:

**(1)** The overall hysteresis error reported by BFEE2 should be less than 10 kcal·mol^−1^. This is the predicted error reported by BFEE2 protocol based on the hysteresis of backward and forward transformations throughout the entire thermodynamic cycle of each replicate [32]. It provides an opportunity to check if a particular replicate has converged enough to provide informative free energy estimates.

**(2)** The free energy contribution of the restraints in the ligand–RNA complex arm should be in a similar range to the calculations done for the other ligands. For this criteria, we check that free energy contributions of restraints calculated in step 1 in thermodynamic cycle (Fig. 2b) is less than 2.5 times the median value for the ΔG calculated from the pool of all replicates of all the ligands, in each condition. Replicates with extreme deviations in restraint contributions were rejected and replaced with a new replicate. An example case is shown in Fig S4b for Method 2, where median of ΔG of restraint contributions in calculateed step 1 was around -2.5 kcal/mol for all replicates and two replicates with much larger magnitude were marked and rejected (shown in grey). An extremely high restraint contribution indicates instability in the ligand pose. In almost all cases, when this red flag was observed, it coincided with large RMSD changes for ligand and RNA being observed in step 1, when restraints were gradually being turned off. RMSD plots of rejected replicates can be seen in Fig. S24.

If either of these conditions were not met for any replicate, we excluded it from our calculation and we ran another independent replicate as a replacement. RMSD plots of accepted replicates are shown in Fig S12 to Fig S23.

### 2.7 RNA-backbone restrained simulations

In some of our simulations, we applied an RMSD restraint to the RNA backbone heavy atoms (atom names: C3’, C4’, C5’, O3’, O5’, OP1, OP2, and P), to retain the RNA backbone conformation observed in the NMR structure [31]. In these simulations, the initial equilibration was performed in two steps as shown in (Fig. 2c): **(1)** 5,000 steps of energy minimization, followed by 4 ns of restrained equilibration by applying harmonic positional restraints (k = 5 kcal·mol^−1^·Å^−2^) to all RNA and ligand heavy atoms as well as the Mg^2+^ ions if present in the system; **(2)** 100 ns of equilibrium simulation with RMSD restraints on the RNA backbone heavy atoms (k = 10 kcal·mol^−1^·Å^−2^), using the NMR structure as the reference.

The last frame from step 2 was used to set up the free energy simulations using BFEE2. Then, in the simulations of the bound system, an RMSD restraint on the RNA backbone heavy atoms (k = 10 kcal·mol^−1^·Å^−2^) was added on top of the other 7 restraints described in section 2.4, by editing the input colvar files (‘000_eq/colvars.in’, ‘001_Molecule-Bound/ colvars.in’, ‘002_RestraintBound/colvars_backward.in’, and ‘002_RestraintBound/colvars_forward.in’). In these simulations, the backbone RMSD restraint was applied with respect to the last frame of step 2 of the initial equilibration. To account for the contribution of the restraints on the RNA backbone atoms, we used the modified thermodynamic cycle shown in Fig. 2d. In this thermodynamic cycle, there is a need to calculate an extra step (step 5) in which we apply the RMSD backbone restraints on the RNA aptamer in the absence of any bound ligand and calculate its energetic contributions. These calculations were carried out using TI simulations in a bidirectional manner, each direction simulated for 40 ns in total, during which the RMSD backbone restraints were gradually turned on and off. We repeated this calculation on the RNA-only system three times and used its average value to account for the contribution in the final binding free energy values for all the compounds. To calculate free energy estimates from simulations with RNA backbone restraints, we follow the BFEE2 post-treatment procedure [56] with the additional change in postTreatment.py script to accept eight collective variables, instead of the original seven ligand-only restraints.

The step-by-step protocol for running free energy calculations with RNA backbone restraint and deviations from the original BFEE2 protocol were delineated in SI Section 8.4.

### 2.8 MM-GBSA Calculations

We used the MM-GBSA (Molecular Mechanics Combined with Generalized Born and Surface-Area Solvation) method [58–60] to estimate the binding free energy of six different ligands, to the RNA aptamer from their equilibrated trajectories.

The binding free energy in solution can be estimated using

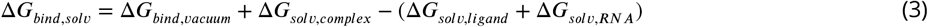

where Δ*G*_*bind,vacuum*_ is the binding free energy in vacuum, Δ*G*_*solv,complex*_ is the solvation free energy of the RNA complex in solution, Δ*G*_*solv,ligand*_ is the solvation free energy of individual ligand in solution, and Δ*G*_*solv,RNA*_ is the solvation free energy of the RNA in solution. Solvation free energies involve electrostatic and hydrophobic contributions where the hydrophobic contribution is an empirical value while the electrostatic contribution is estimated using the Generalized Born equation for each of the above three states. We can estimate Δ*G*_*bind,vacuum*_ using the following equation:

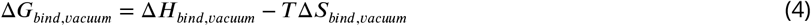

where Δ*H*_*bind,vacuum*_ is the enthalpy of binding, T is temperature and Δ*S*_*bind,vacuum*_ is the entropy of binding which is estimated using the Normal Mode Analysis.

We used the three replicates of 100 ns pre-BFEE2 equilibration simulations collected for each ligand system for MM-GBSA calculations. 100 frames were extracted from each trajectory, uniformly distributed in time, and re-imaged to center RNA in the middle of the periodic box, using the LOOS analysis package [61, 62]. For each ligand, free energy estimates were obtained from three replicate equilibrium trajectories, using the MMPBSA.py script [60] from Amber-Tools version 23 [63] to perform MM-GBSA calculations. Final estimates were reported as the mean and standard deviation of three replicates for each ligand.

We performed the MM-GBSA calculations in two ways. The first MM-GBSA calculation (Method 20 in Table 1) was performed from equilibrium trajectories with 55 mM NaCl with two Mg^2+^ originally prepared for Method 2. The second MM-GBSA calculation (Method 21) was performed from equilibrium trajectories with 55 mM NaCl without Mg^2+^, originally prepared for Method 1. In both cases, TIP3P water model and GAFF2 small molecule force field were used.

Detailed protocol for MM-GBSA calculations is provided in SI Section 8.6.

### 2.9 Analysis of Performance

For each ligand and explored method, we ran three replicate calculations and reported the average and standard deviation of calculated binding free energies. We compared the binding free energy of the compounds with their corresponding experimental measurements and evaluated the results using several different statistical metrics, including root mean square error (RMSE), mean absolute error (MAE), Pearson’s correlation coefficient (*r*), Kendall’s rank correlation coefficient (*τ*), and Spearman’s rank correlation coefficient (*ρ*).

To assess the stability of the RNA aptamer and the bound ligand, the RMSD was calculated by first aligning the RNA backbone heavy atoms (atom names: C3’, C4’, C5’, O3’, O5’, OP1, OP2, and P) to the experimental NMR structure of the RNA aptamer. We plotted the time evolution of both RMSD of the RNA backbone heavy atoms and ligand heavy atoms through all stages of the protocol (See Figure S12 to S24. This approach allowed us to catch lower quality replicates in which ligand pose was unstable during the FEP step for capturing complex interactions or during the TI step for capturing restraint contributions (steps 3 and 4 of the thermodynamic cycle as shown in Fig 2). Some examples of these low-quality replicates are shown in Figure S24.

We calculated RNA heavy atom radius of gyration observed in the aggregated trajectory, i.e. pre-BFEE2 equilibration and the trajectories of the alchemical protocol, to compare the effect of including different number of Mg^2+^ ions and RNA backbone restraints on the RNA dynamics (Figure S5-S6).

To check the health of the alchemical protocol, we analyzed the overlap between potential energy distributions of backward and forward calculations at every *λ*-window. To do so, we used ParseFEP [64] to get the probability distributions of the potential energy difference (Δ*U*) values in each lambda-window for backward and forward transformations. Then, we adopted Kullback-Leibler Divergence (KL-Divergence) calculations to quantify the difference between the forward and backward potential energy distributions for easier visualization of problematic transitions as shown in Figure S7. A detailed protocol of KL-Divergence calculations for potential energy overlap assessment can be found in SI Section 8.7. We also monitored the overlap in potential-energy distributions in this exercise of doubling the sampling. Fig. S8-S11 show examples of KL-Divergence plots. Green bars indicate pairs of *λ* states with sufficient overlap in potential free energy distributions and red bars indicate alchemical transformations with relatively lower overlap, ie. higher divergence. This visualization allowed a quick look into how alchemical protocol changes were impacting the energetic overlap of sampled states.

We investigated the differences in how different monovalent cations (Na^+^ and K^+^) interact with RNA. We plotted the radial distribution function (RDF) of monovalent cations to the two oxygen atoms of the RNA backbone phosphates to understand their spatial distribution around the phosphate backbone. To determine the high occupancy sites for monovalent cations we calculated the density of cations using the VolMap plug-in in VMD [47]. The cation occupation probability densities were averaged throughout the 100 ns pre-BFEE2 equilibration trajectory. We visualized cation sites with relatively higher occupancy by depicting isovalue surfaces at 0.03. For these visualizations, both K^+^ and Na^+^ van der Waals radii were set to 1 Å before calculating occupation probability densities to remove the effect of the cation size on the depiction of high occupancy sites.

### 2.10 Computing resources

All calculations for this study were performed on Amazon Web Services. The pre-BFEE2 equilibration took about 13 hours on p3.2xlarge instances with NVIDIA Tesla V100 GPUs and Intel Xeon Scalable Processor (Broadwell E5-2686 v4), with the cost of $3.06 /hour. For the BFEE2 simulations, g5.8xlarge instances were used with NVIDIA A10G Tensor Core GPUs and 2nd generation AMD EPYC processors (AMD EPYC 7R32). All BFEE2 steps, except for the systems with OPC water, were completed in about 74 hrs with the cost of $1.624 /hour for each replicate of each ligand. With 100 ns pre-equilibration and alchemical protocol of the 40 *λ*-windows and 1 ns/window that relies on three replicate simulations to estimate the free energy of each ligand full calculation cost per ligand was close to $468 for each investigated simulation condition.

For systems with the OPC water model, the pre-BFEE2 equilibration was performed using CUDA-accelerated NAMD2.14, instead of NAMD3. Using g5.8xlarge instances, this step was completed in about 33 hours. For steps 1 and 4 of the thermodynamic cycle were done with CUDA-accelerated NAMD2.14, however, steps 2 and 3 are done with NAMD2.14.

The total time for completion of the BFEE2 steps on g5.8xlarge instances was around 192 hours.

## 3 Results and discussion

In this study, we aimed to investigate the performance of alchemical free energy methods in predicting absolute binding free energy of small molecules to RNA. To achieve this, we studied the binding affinity of theophylline and five of its analogs to the RNA aptamer. We examined the impact of various simulation setup decisions, including different salt conditions, Mg^2+^ placements, water models, force fields describing the interactions of ligands, simulation time, and lambda schedules, on the results of our calculations. Table 1 summarizes the various conditions studied in this study.

### 3.1 Mg^2+^ placement is a critical decision for binding free energy calculations

One of the major challenges in calculating the binding free energies of ligands to RNA targets is the inherent flexibility of RNA. The secondary and tertiary structure of RNA is highly dependent on the presence of divalent cations, such as Mg^2+^ [65–67]. The Mg^2+^ ions stabilize the structure of RNA by binding to specific parts of the RNA backbone, but the exact locations of these binding sites are not always known for different RNA targets. For the theophylline binding aptamer, three Mg^2+^ ions are suggested to bind the aptamer’s backbone [22, 44, 45]. 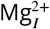 is the closest to the ligand binding site out of the three, and is in the U-turn formed by C22, U23, and U24, whereas 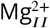 coordinates with the phosphate groups of G14-A16, and 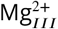 is located at the G2:U32 wobble pair region in the lower stem of the aptamer [22] (Fig. 1). To evaluate the effect of Mg^2+^ placement on binding free energy calculations, we examined systems with zero, two, and three Mg^2+^ ions. For the two Mg^2+^ system, we included 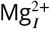 and 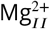, as shown in Fig. 1.

In all three cases, we added 55 mM NaCl to each system and calculated the binding free energy of theophylline and its five analogs to the RNA aptamer. We replicated each system three times and reported the average binding free energy from these replicas, with the error bars indicating the standard deviation in Fig. 3a-c. Without Mg^2+^, the standard deviation between replicas for each compound is significantly larger than in the case of two and three Mg^2+^ systems (Fig. 3a-c). This could be related to the role of Mg^2+^ ions in stabilizing the RNA structure.

**Figure 3.**
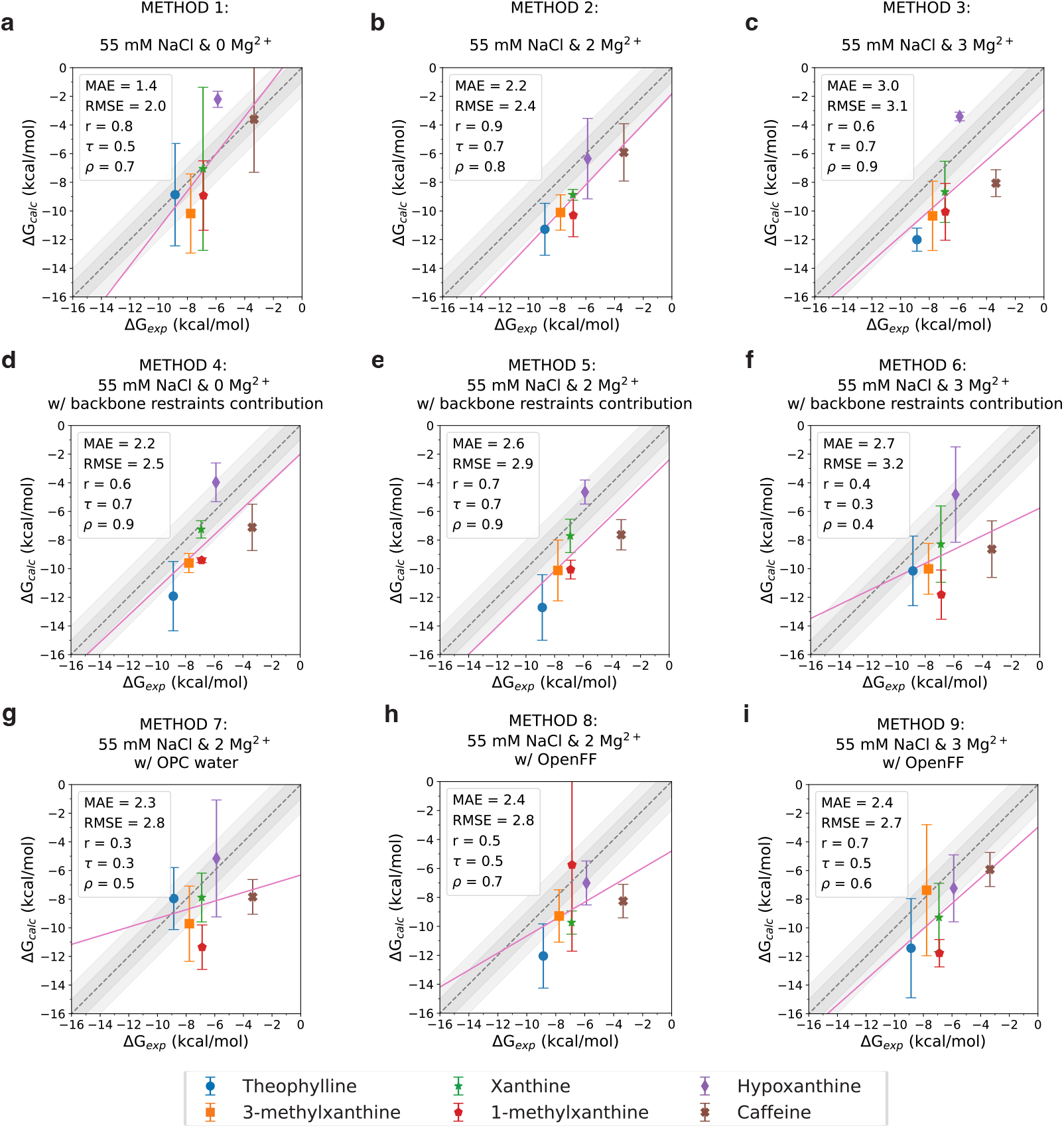
Experimental vs. calculated binding free energies. Each compound is represented with a different color, with each data point representing the average of the three accepted replicate simulations and the error bars showing the standard deviation among the three replicates. The identity line is shown as a dashed line, with 1 and 2 kcal/mol deviations shown in shades of gray. MAE (kcal/mol), RMSE (kcal/mol), Pearson’s correlation coefficient (r), Spearman’s rank correlation coefficient (*ρ*), and Kendall’s rank correlation coefficient (*τ*), are listed in the legends. **(a), (b)**, and **(c)** Binding free energies for theophylline and its analogs for systems with 55 mM NaCl and TIP3P water model and zero, two, and three Mg^2+^ ions, respectively. **(d), (e)**, and **(f)** Systems with RNA backbone restraints and 55 mM NaCl and TIP3P water model and zero, two, and three Mg^2+^ ions, respectively. **(g)** Similar to **(b)** but with OPC water model instead of TIP3P. **(h)** and **(i)** Similar to **(b)** and **(c)**, respectively, but with the ligand’s force field described by OpenFF 2.0.0 Sage [42, 43].

The two Mg^2+^ system (Method 2) has the best overall performance in terms of correlation with *r* = 0.9, and *τ* = 0.7, and *ρ* = 0.8. In this case, the calculated binding affinities are overestimated by ≈ 2.2 kcal·mol^−1^, as indicated in Fig. 3b. The addition of the third 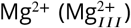, negatively impacts the results, increasing MAE to 3.0 kcal·mol^−1^ (Method 3). This is possibly because of the mischaracterization of the 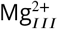 placement on the RNA aptamer. Consistent with this observation, a recent MD study also reported the instability of the 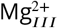, and its dissociation within a few nanoseconds of simulation [28].

The sensitivity of prediction performance to Mg^2+^ ion placement is a challenge for setting up free energy calculations for prospective predictions. In this study, we learned which Mg^2+^ ion positions to consider from prior studies. Still, among different combinations, we were only able to judge which placement was better based on prediction accuracy relying on experimental affinity values. The decision of Mg^2+^ placement would only get more difficult for more complex and less studied RNA targets, especially if experimental evidence for high-affinity Mg^2+^ binding sites is not available. The modeling decisions around Mg^2+^ ions would be especially challenging for prospective prediction of RNA-ligand interactions.

### 3.2 Restraining RNA backbone is a safe alternative to dificult Mg^2+^ placement decisions for restricting conformational space

To address the flexibility of RNA and lack of information on Mg^2+^ binding sites, we explored the impact of applying restraints to limit conformational changes in RNA. We applied an RMSD restraint to the RNA backbone heavy atoms (k = 10 kcal·mol^−1^·Å^−2^) and added one more step to calculate the contribution of this target RMSD restraint, as shown in step 5 of the thermodynamic cycle in Fig. 2d. We tested the application of the RMSD backbone restraints with zero, two, and three Mg^2+^ ion-containing systems as well (Methods 4, 5, and 6, respectively). Our results indicated that even if we did not include Mg^2+^ in our system, the application of the RMSD backbone restraints improved the convergence of the binding free energy calculations. The standard deviation between replicas decreased with the use of RMSD restraints compared to the systems without restraints, Fig. 3a,d. Having Mg^2+^ in the system while applying the backbone restraints seems to not impact the results, as long as the Mg^2+^ are placed in the correct position, which is the case for the two Mg^2+^ system, Fig. 3e (Method 5). Surprisingly, even with the RNA backbone restraints, the addition of the 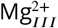 seems to negatively impact the results, Fig. 3f (Method 6). Overall our results show the promise of using restraints on the RNA structure as a solution for cases where there is a reliable experimental 3D structure of the RNA target, but no experimental guidance on the Mg^2+^ binding sites.

### 3.3 Combination of Amber OL3 with TIP3P water and GAFF2 ligand parameters performed better than tests with OPC water or OpenFF Sage force field

To evaluate the impact of the force field for modeling small molecules, in addition to GAFF2 [39], we used OpenFF 2.0.0 Sage [42, 43] to describe the ligands in our systems. We tested the Sage force field in two conditions: 55 mM NaCl, without backbone restraints, with two or three Mg^2+^ ions (Method 8 and 9, respectively). For both the MAE values were found to be slightly lower with Sage (2.4 kcal/mol instead of 3.0 kcal/mol), but we observed a reduction from 0.7 to 0.5 for Kendall’s rank correlation coefficient. Overall, the Sage force field did not provide any improvement over results achieved with GAFF2 (Method 2), as shown in Fig. 3c,g. Therefore, we decided not to pursue further exploration of the Sage force field in other conditions.

Although this force field has many reported successes in the areas of predicting solvation free energies and proteinligand binding affinities, for this model system we did not observe an improved performance with Sage over GAFF2. Boothroyd et al. reported that Sage was developed to be compatible with Amber-family protein force fields [68], but it may be less compatible with the Amber RNA force field. It is known that modeling ligand-RNA interactions has not been a consideration for the development of Sage. This can be remedied in the future releases of OpenFF force field or could be addressed by co-development of small molecule and RNA force fields in the future.

Most of our calculations were performed using the TIP3P water model, as the Amber OL3 force field for RNA was developed with TIP3P [38]. We specifically wanted to test the effect of the OPC water model in Method 7 with 55 mM NaCl and two Mg^2+^ ions, Fig. 3b,h. OPC was chosen as the alternative as this model was reported to more accurately capture bulk properties of water [41] and in recent studies OPC water model was reported to capture experimental RNA structures better especially when used with additional modifications to Amber OL3 force field, such as adjustment to van der Waals radii of phosphate oxygen atoms [69]. We did not explore these modifications and used Amber OL3 force field as is in this study. With OPC (Method 7), Kendall’s Tau dropped to 0.3 from 0.7, which was achieved with equivalent simulations with TIP3P (Method 2). Meanwhile, the MAE with OPC remained relatively similar to that of the system with the TIP3P water model. Overall, we did not observed any benefit for using OPC instead of TIP3P water when RNA is modeled with Amber OL3 force field. Additionally, OPC water reduced the speed of calculations significantly, especially due to its incompatibility with NAMD GPU code. We had to run these calculations using only CPUs.

### 3.4 Mimicking the buffer conditions of the binding afinity experiments improved the accuracy of free energy calculations

We observed that RNA-ligand free energy calculations were sensitive to both ionic strength and salt identity of the aqueous environment. Simulation conditions with 55 mM NaCl represent our closest approximation of the experimental conditions in terms of cation identity and ionic strength.

To compare the effect of monovalent cation choice, we also simulated systems with 55 mM KCl containing two and three Mg^2+^ ions (Method 10 and 11, respectively). Compared to the systems with NaCl with matching ionic strength, surprisingly, the binding free energies results with KCl were farther from the experimental values (Fig. 4a, b). For a deeper dive into the differences between how these cations interact with RNA please see Section 3.5.

**Figure 4.**
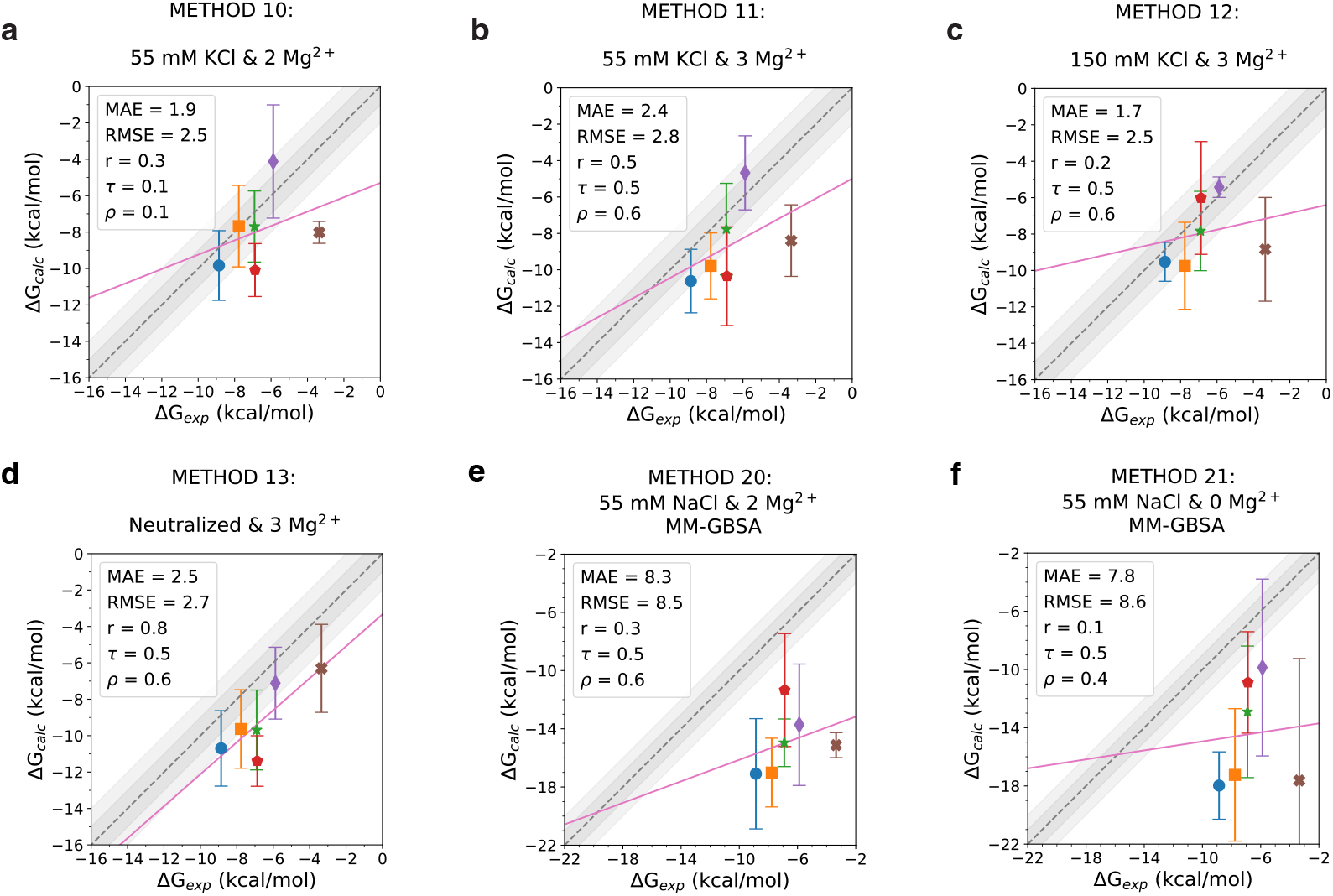
Effects of salt condition on binding free energy calculations and comparison to MM-GBSA calculations. Experimental vs. calculated binding free energies for different conditions: **(a)** 55 mM KCl and two Mg^2+^, **(b)** 55 mM KCl and three Mg^2+^, **(c)** 150 mM KCl and three Mg^2+^, **(d)** Neutralized only system with three Mg^2+^ ions without any additional salt beyond neutralization, **(e)** MM-GBSA calculation using 55 mM KCl and two Mg^2+^ system, and **(f)** MM-GBSA calculation using 55 mM KCl and zero Mg^2+^ system. Each compound is represented with a different color, with each data point representing the average of the three accepted replicas and the error bars showing the standard deviation among the three replicate simulations. Refer to the legend of Fig 3 for ligand colors. The identity line is shown as a dashed line, with 1 and 2 kcal/mol deviations shown in shades of gray. Mean absolute error (MAE, kcal/mol), root mean square error (RMSE, kcal/mol), Pearson’s correlation coefficient (*r*), Spearman’s rank correlation coefficient (*ρ*), and Kendall’s rank correlation coefficient (*τ*) are listed in the legends.

We explored two extreme conditions to evaluate how important it is to approximate the ionic strength of the experimental buffer environment in free energy calculations: In Method 13, to show the detrimental effect of ignoring ionic strength adjustment, we only neutralized the system with Na^+^ ions and structural Mg^2+^ ions and did not add more salt (Fig. 4d). Method 13 has slightly lower rank correlation coefficients (Tau = 0.5, rho = 0.6) compared to the equivalent condition with 55 mM NaCl and three Mg^2+^ (Method 2, Tau = 0.7, rho = 0.9). Increasing the salt concentration to 150 mM KCl with two Mg^2+^ ions (Method 12) also does not improve the results (Fig. 4c). Based on these observations we think that it is more important to model salt conditions correctly for free energy calculations of RNA targets compared to protein targets, since for protein-ligand free energy calculations common practice is just including neutralizing counter ions, not necessarily approximating the experimental ionic strength. Due to RNA’s highly negatively charged backbone, ionic strength and the shielding effect of salt molecules are expected to play a bigger role in RNA conformation.

### 3.5 Differences in spatial distribution of monovalent cations around RNA can be the reason for Differences in free energy calculations

Cation size-dependent stabilization of RNA structures has been shown in single-molecule optical and magnetic tweezer experiments, nuclear magnetic resonance, and gel electrophoretic studies. In these experiments, RNA stability is higher in NaCl solution compared to KCl solution [70]. Hence, we investigated the binding preference of Na^+^ and K^+^ to the RNA surface in NaCl and KCl solutions. We first looked at the radial distribution function (RDF) of Na^+^ and K^+^ with respect to the phosphate groups of the RNA, Fig. 5a. Our analysis suggests that Na^+^ ions condense more onto the RNA. The contact peak center for K^+^ ions is shifted 0.5 Å away according to the RDF plot compared to Na^+^, which match the difference in van der Waals radius of two ions. But based on the area of the first RDF peak, more Na^+^ ions were found to be in contact with RNA backbone compared to K^+^ when modeled with TIP3P water. This is in line with a recent study that reported smaller hydrated sodium ions condensing more around the phosphate groups, leading to a reduction of electrostatic repulsion and enhancing RNA stability.

**Figure 5.**
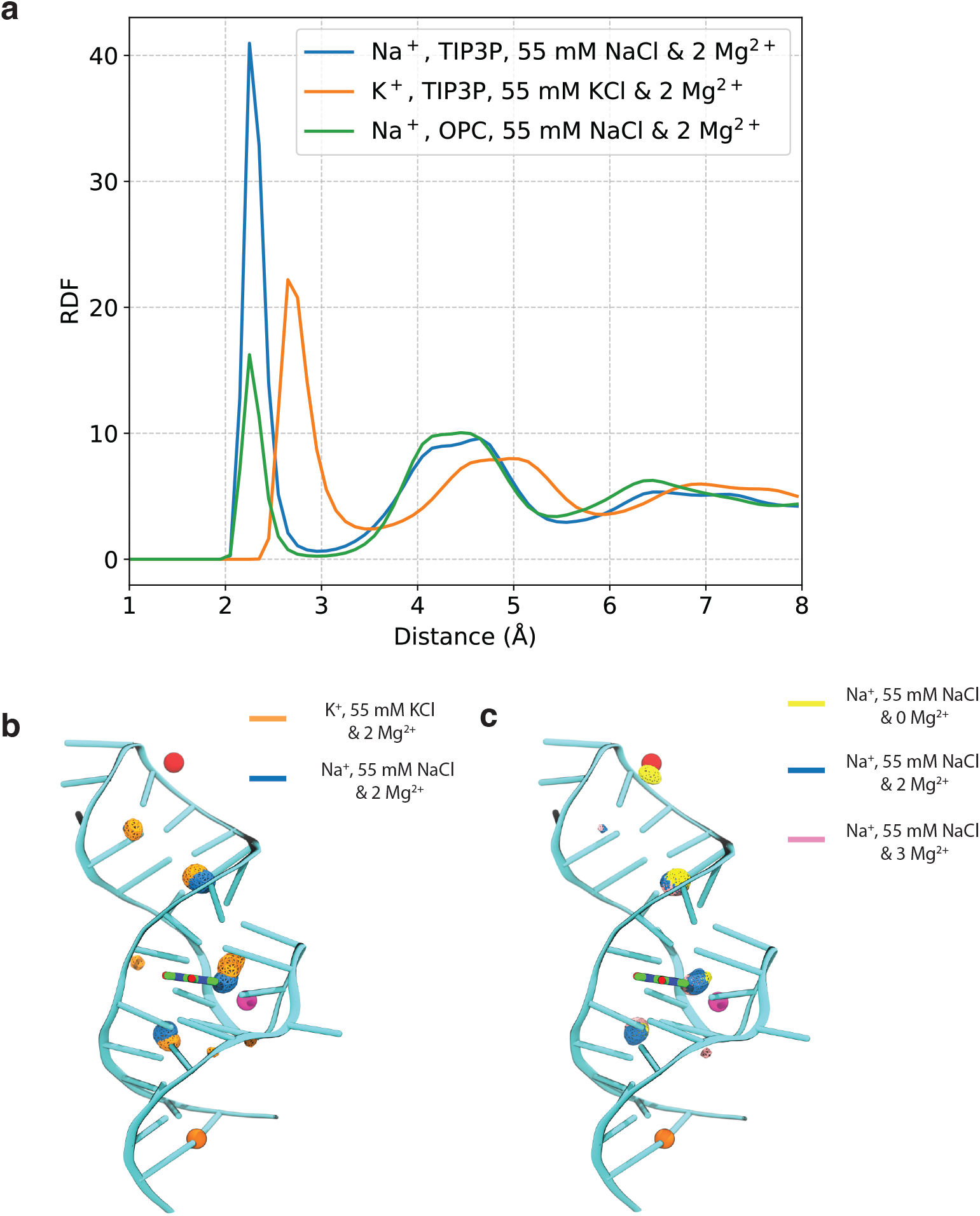
Distribution of monovalent cations around RNA. **(a)** Radial distribution function (RDF) plot of monovalent cations with respect to RNA backbone on systems with 55 mM NaCl and two Mg^2+^ with TIP3P water model (blue line), 55 mM KCl and two Mg^2+^ ions with TIP3P water model (orange line), and 55 mM NaCl and two Mg^2+^ with OPC water model (green line). The RDF calculations were performed calculating the distance of cations from the two oxygen atoms of the RNA backbone phosphates. **(b)** Density of cations, K^+^ and Na^+^ averaged throughout the 100 ns pre-BFEE2 equilibration trajectory, calculated using Volmap tool in VMD. All densities are shown at isovalue 0.03. Both K^+^ and Na^+^ Van der Waals radii were set to 1 Å before the calculations. **(b)** Density of Na^+^ ions in 55 mM NaCl with zero, two, and three Mg^2+^ systems shown in yellow, blue, and pink, respectively. In the zero Mg^2+^ system, there is an extra cation density where 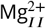 resides in the other two systems.

Additionally, we calculated the density of the cations and visualized the high density regions around the RNA using aggregated trajectories of all ligands, Fig. 5b. We observed that the highest occupancy sites of Na^+^ and K^+^ ions are typically adjacent but show some differences. Most notably, the location of the cation binding site between the small molecule and the 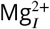 is slightly shifted in simulations with K^+^ relative to ones with Na^+^. Positioning of monovalent cations, especially near the binding site may be a factor affecting the free energy calculations.

We also evaluated the changes in monovalent cation binding sites between simulation conditions with zero, two, or three Mg^2+^ ions, as shown in Fig 5c. The most interesting observation was that when 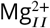 was omitted from simulations, a high occupancy Na^+^ site appeared in that position. However, this behavior was not observed for the 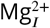 and 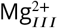 positions.

### 3.6 Increasing the number of *λ* windows provides a slight improvement while increasing sampling time per window can be detrimental

Based on the results of the individual steps in the thermodynamic cycle, step 3 which involves complexation free energy calculations in the binding site, shows the highest variance between independent replicas that are simulated (see Fig. S4a and S4e, in comparison to other steps). To check if increasing the sampling in this step can improve the results, we doubled the sampling of step 3 (Fig. 2b), in two different ways: by doubling the number of *λ* windows (80 windows, 1 ns/window; Method 16) and by doubling the sampling time per *λ* window (40 windows, 2 ns/window; Method 17), see Fig. S3. We tested out doubling the sampling in two conditions: 55 mM NaCl with two Mg^2+^ ions, and 55 mM NaCl without Mg^2+^ and with RNA backbone restraints (Methods 18 and 19). Compared to our base calculations (40 win, 1 ns/win; Methods 2 and 5), using 2 ns/win in both simulated conditions (Method 15 and 17), not only does not improve the results, but also slightly underperforms in both simulation conditions (Fig. S3b, d and Fig. 3b, d). The reason longer simulation time per window causes deterioration in prediction accuracy could be due to the flexible nature of RNA and the deficiency of the force field. We hypothesize that increasing the sampling time diminishes the result quality because the RNA conformation drift away from the experimental structure. Using 80 *λ* windows of 1 ns, on the other hand, shows similar rank ordering performance in both conditions (Methods 14 and 16, compared to Methods 2 and 5). Moreover, in the case with the RNA backbone restraints, Method 18 (80 windows of 1 ns) shows slight improvement in MAE and RMSE compared to the base calculations (Method 4).

Considering the cost-benefit ratio, we believe our initial sampling scheme was already sufficient, and further increasing the sampling did not lead to significant improvements. These findings suggest the importance of careful evaluation of the sampling efficiency in alchemical free energy calculations to optimize the computational resources and improve the accuracy of the calculations.

### 3.7 Free energy calculations provided better predictive power compared to MM-GBSA approach at a greater computational cost

Running free energy protocol with 40 *λ* windows, 1 ns/window and three replicates for each ligand has significant calculation cost, roughly $468 per ligand including the pre-BFEE2 equilibration runs using the AWS instances described in Section 2.10. The cost of these calculations poses a limitation on the number of RNA-ligand systems that can be routinely evaluated. We estimate that for a typical discovery project, it would be only feasible to evaluate tens of ligands with this approach. With this in mind, we were motivated to explore how the performance of absolute free energy calculations would compare to a less expensive approach: MM-GBSA calculation to estimate binding free energy repurposing the 100 ns pre-BFEE2 equilibration runs.

The accuracy of absolute binding free energy predictions was significantly lower with MM-GBSA calculations. MAE of 8.3 and 7.8 kcal/mol was observed for MM-GBSA calculations in Method 20 and 21, respectively. In both cases, free energy estimates from MM-GBSA calculations significantly overestimated binding affinities for all ligands. The overestimation of binding affinities by MM-GBSA is expected to get larger with larger ligands.

For the ranking performance, the free energy calculations (Method 2, 4, and 5) were more successful than MM-GBSA approach. This result was expected since MM-GBSA calculations were not predicted to do well in the polar and water-exposed binding sites such as binding sites in RNA targets [58]. Nevertheless, MM-GBSA calculations demonstrated some ability to rank ligands (Kendall’s Tau of 0.5) and calculations cost a quarter of the cost of free energy calculations. Considering the lower cost, the MM-GBSA approach may still be preferable as a first-pass enrichment method when it is necessary to evaluate a larger pool of ligands.

### 3.8 Comparison of performance to other studies

There were a few prior studies which studied free energy calculations on the theophyllin-binding RNA aptamer: The first study by Gouda et al. provides a comparison of the molecular mechanics Poisson–Boltzmann surface area (MM-PBSA) and Thermodynamic Integration (TI) approaches for predicting relative binding affinities of six ligands, studied in our paper [22]. MM-PBSA calculations based on computational mutagenesis of the ligand on snapshots from a single MD trajectory approach achieved an impressive r^2^ of 0.82 (Pearson’s r of 0.90), given the simplicity of the approach. TI calculations were carried out to calculate relative free energy differences (ΔΔ*G*). In contrast to our approach, the alchemical transformation was sampled much more narrowly with 101 windows and 3 ps equilibration and 3 ps data collection, however, authors reported successful results with r^2^ of 0.98 (r of 0.99). By contrast, we had to run the absolute free energy calculations much longer with 40 windows 0.2 ns equilibration and 0.8 ns data collection to obtain our best results with Pearson’s r of 0.9 with Method 2. This highlights the advantage of the relative free energy calculations over absolute whenever ligands are congeneric with small perturbations and expected to occupy similar binding poses. For our study design, we had chosen to pursue absolute free energy calculations despite the expected sampling challenges and increased cost, due to the attractiveness of the broader application area of diverse ligands and poses.

Tanida et al’s paper, published in 2007, reports absolute binding free energy prediction on the same model system with the approach of estimating nonequilibrium work using MC-CAFEE protocol [29]. They achieved similar performance and calculation efficiency to the TI approach presented by Gouda et al. [22] for estimating relative affinities of ligands (Pearson’s r of 0.99). However, they also reported a constant bias in absolute free energy estimates: Δ*G*_*calc*_ values were all 7 kcal/mol lower compared to the experimental values for all ligands. In comparison, we were able to achieve a MAE of 2.2 for our best methods (Method 2 with three Mg+2 ions, Method 4 with backbone restraints, and Method 16 with doubled *λ* schedule), although our computational cost was significantly higher. Tanida et al. reported that the hypoxanthine system has the highest deviation from the linear relationship between predicted and calculated free energies (1.5 kcal/mol) and suggested that the binding site of hypoxanthine could be different than others. Similarly in our hands, hypoxanthine estimates have the highest deviation from linearity in most of our tested methods (see correlation plots of Method 1-6 in Fig 3). We observed that hypoxanthine affinity was systematically underestimated, which suggests that perhaps it was not modeled in the right pose or binding site. In both of these early studies, calculations were run only once, so we were not able to learn about the prediction uncertainty and the reproducibility of estimates.

In 2019 Tanida et al. published a second paper revisiting the free energy calculation of theophylline-RNA aptamer complex, this time focusing only on theophylline and not the other analogs [28]. The goal of this paper was demonstrating how both binding poses and binding affinities can be predicted for an RNA ligand. In this paper, they first discover different binding poses of theophylline in the experimentally known binding site with metadynamics, compute free energy of binding for six different poses separately, and then estimate an overall free energy of binding with contributions from all poses. For the alchemical protocol they used 6 *λ* and 14*λ* windows for gradually turning off Coloumb and Lennard-Jones interactions sequentially with 2 ns equilibration and 6 ns production run per window. This exhaustive protocol achieved a very accurate predicted binding free energy for theophylline, exactly matching the experimental binding affinity of -8.9 kcal/mol. It would be interesting to explore how this approach would perform for ranking the binding affinities of theophylline analogs. The metadynamics-based determination of binding poses could be too challenging for applying this method to tens of ligands at once, due to the need to engineer useful collective variables and monitor convergence. We suspect docking-based pose predictions would be more practical to implement for automated protocols designed to compare many diverse ligands, although, errors in the initial pose might reduce the quality of the results. An alternative approach could be resorting to nonequilibrium candidate Monte Carlo steps to improve sampling of ligand binding modes [71]. Binding pose prediction and its effect on binding affinity estimates was beyond the scope of our paper. For learning more about recent improvements and remaining challenges of RNA docking we recommend these two reviews [72, 73].

Both Tanida et al. studies mentioned above used Amber force fields for RNA (ff99 and ff14SB), GAFF for ligand parameters, and TIP3P for the water model. Our observations also support their choice of ligand force field and water models. None of these three studies have investigated the effect of simulation setup decisions such as including Mg+2 ions or approximating the ionic strength of the experimental affinity measurements with NaCl or KCl salts. So we hope our systematic comparison will provide useful guidance for future applications of alchemical free energy methods to RNA-ligand systems.

### 3.9 Limitations of this study and future work

It is important to acknowledge that only one RNA target has been used in this study and six congeneric ligands. The suggestions we highlighted and found useful in this study would surely benefit from being tested with a diverse set of RNA targets and ligands before being considered as general rules for modeling RNA-ligand binding. Because absolute free energy calculations with RNA targets are largely unexplored territory, we started with baby steps before attempting to run with drug-like systems and therapeutic targets.

We chose the theophylline binding RNA aptamer as the target to study because of the availability of an experimental 3D structure with at least one ligand and its simple hairpin structure. Larger and more flexible RNA structures can pose difficulties for free energy calculations in terms of convergence and calculation costs. The existance of prior studies on Mg^+2^ ion placement was also helpful to us, guiding the possible positions we explored in this study. Mg^+2^ placement decision is expected to be much harder for RNA systems that will be modeled for the first time.

The simplicity of its ligands was one of the most important features that made us prefer this model system. The six theophylline analogs are all neutral and rigid small molecules which is admittedly less challenging for free energy calculations compared to a typical drug-like molecule. Therefore, the prediction performance we observed in this model system can be optimistic for ligands with higher complexity. Obvious next steps to explore include checking how the performance evolves with increasingly complex ligands such as charged ligands or ligands with rotatable bonds.

The dynamic range of ligand affinities of the current set poses a limitation for being able to distinguish performance differences. The free energy difference between the highest and lowest affinity ligands in this dataset is 5.51 kcal/mol, which is just enough be able to distinguish broadly good performance from bad performance. A data set with a larger dynamic range of affinities would have allowed us to have a more detailed comparison between performance levels, but this was the broadest ligand range we could find in any RNA system we considered.

Currently, the ability to expand this study to a diverse set of targets and small molecules is limited by the availability of RNA-ligand benchmark sets and the cost of calculations. There are a number of RNA-ligand complex structures in the PDB Database based on X-ray or NMR measurements, however, it is much harder to find RNA targets that have both experimental 3D structures and a series of known ligands with a sufficiently large range of experimentally determined affinities and known binding sites. The field of RNA free energy calculations can benefit from the construction of broader benchmark datasets with large target and ligand diversity and wide range of affinities.

## 4 Conclusion

In this study, we evaluated the performance of absolute free energy calculations for predicting the affinity of six theophylline analogs to theophylline-binding RNA aptamer. The goal was to understand the prediction performance of free energy calculations for a simple RNA-ligand complex system. In contrast to protein targets, running free energy calculations for RNA targets is not common and we wanted to learn about specific modeling considerations that impact the success of RNA binding predictions. We used BFEE2 to automate the execution of absolute binding free energy calculations. To our knowledge, this is the first application of BFEE2 to free energy calculations of an RNA target. Systematic exploration of various modeling decisions about Mg^2+^, salt conditions, backbone restraints, ligand force fields, and water models led to valuable insights. We observed that Mg^2+^ placement has a significant effect on the performance. Ignoring Mg^2+^ ions or adding extra ions was detrimental to the predictive performance. We showed that the prediction accuracy of the best Mg^+2^ placement can be recapitulated by implementing RMSD-based RNA backbone restraints. For the studied system, restraining the RNA backbone throughout the free energy calculation turned out to be a safer alternative to Mg^+2^ placement for managing RNA flexibility. Detailed modifications to BFEE2 protocol were provided for implementing backbone restraints for the target molecule and to account for it correctly in the thermodynamic cycle. We also observed that RNA free energy calculations were sensitive to both ionic strength and the identity of monovalent cation while mimicking experimental buffer conditions led to the best results. Compared to typical protein-ligand free energy calculations, calculated free energies for the RNA target in this study showed much higher variability. Therefore, it was necessary to obtain three independent free energy estimates from three different runs for each ligand and to calculate the final free energy estimate as the mean. We also implemented a blind strategy to flag problematic replicates, using quality criteria obtained from the simulations themselves and not relying on experimental results. Our observations on which simulation setup decisions led to better free energy prediction performance will guide future free energy calculations for RNA targets. However, it is important to acknowledge that only one RNA target has been used in this study and six neutral and rigid small molecule ligands. The highlighted suggestions in this study would surely benefit from being tested with a diverse set of RNA targets and ligands before being considered as general rules for modeling RNA-ligand binding. Currently, the ability to expand this study to a diverse set of targets is limited by the availability of RNA-ligand benchmark datasets, experimental RNA-ligand complex structures, and cost of calculations. Until larger studies can be conducted, we hope that the this systematic study of theophylline aptamer system provides a starting point for scientists using free energy methods to predict RNA binding affinities and help them understand the expected performance levels using a state of the art free energy method.

## Abbreviations

RMSE: Root mean squared error
MAE: Mean absolute error
*τ*: Kendall’s rank correlation coefficient
*ρ*: Spearman’s rank correlation coefficient
r: Pearson correlation coefficient
RDF: Radial distribution function
vdW: van der Waals
RNA: Ribonucleic acid
mRNA: Messenger RNA
RMSD: Root mean squared deviation
BFEE2: Binding Free Energy Estimator 2
CV: Collective Variable

## 5 Code and protocol availability

- Detailed step-by-step simulation protocols for absolute free energy calculations and MM-GBSA calculations are provided in Supplementary Information Section 8.1.
- Input files and analysis scripts are available at https://github.com/modernatx/rna_theophylline_free_energy_calculations.

## 6 Author Contributions

Author contributions are specified below following CRediT Taxonomy.

Conceptualization, MIB; Methodology, AR, MIB, AG, SS; Software, AR; Formal Analysis, AR, MIB; Investigation, AR, MIB, SS, FCP; Data Curation, AR, MIB; Writing-Original Draft, AR, MIB; Writing - Review and Editing, AR, MIB, SS, AG, FCP; Visualization, AR, MIB; Supervision, MIB; Project Administration, MIB; Resources, MIB, FCP; Funding Acquisition, MIB, FCP.

## 7 Acknowledgments

Authors acknowledge support from Moderna Platfrom Research and Discovery Chemistry leadership for resources to conduct this study. We thank Ashlin James Poruthoor for comparing the geometric and alchemical routes of BFEE2 tool on a protein target system and giving us guidance to focus on the more automated alchemical route. His insights were valuable to us for understanding which protocol of BFEE required less manual intervention and more appropriate for dealing with large set of ligands. We are thankful to Semiha Kevser Bali for guidance on adopting the Amber MM-GBSA protocol to RNA targets.

## 8 Disclosures

This study was funded by Moderna, Inc. Mehtap Isık Bennett, Sreyoshi Sur, and Frank. C. Pickard are current employees of Moderna, Inc., and Ali Rasouli is a past employee. They may hold stock/stock options in the company.

## Notes

https://github.com/modernatx/rna_theophylline_free_energy_calculations

### Competing Interest Statement

This study was funded by Moderna, Inc. Mehtap Isik Bennett, Sreyoshi Sur, and Frank. C. Pickard are current employees of Moderna, Inc., and Ali Rasouli was a past employee. They may hold stock/stock options in the company.

