## Supplementary Information Document for "Essential Considerations for Free Energy Calculations of RNA-Small Molecule Complexes: Lessons from Theophylline-Binding RNA Aptamer"

### 8.1 Simulation protocol

In this section, we present the detailed protocol used in this study to prepare and perform alchemical free energy calculations using a single ligand (xanthine) as example. The files referred in the protocol can be accessed from the GitHub repository.

**Description of the directory names**: In the main directory of the repository, you can find this path `‘./fe_calcs_with_0_or_2_mg/6-xanthine/3-55NaCl_Mg/1-40winCmplx_30winLig/1-rep1’`. This path points to the files for the ligand (xanthine), the buffer conditions (55 mM NaCl + 2 $Mg^{2+}$), number of windows used in the alchemical transformation of the RNA-ligand complex (40 windows) and ligand only system (30 windows), and lastly the replica number.

#### 8.1.1 System preparation

**Aim**: Prepare coordinate and topology files, compatible with NAMD MD engine, for a system containing: theophylline-binding RNA aptamer + xanthine bound in the binding pocket + 2 $Mg^{2+}$ ions bound to the RNA backbone, $Mg^{2+}_{I}$ and $Mg^{2+}_{II}$, see section 2.1 + water box + 55 mM NaCl.

1. Go to the system preparation directory:
   `‘cd ./fe_calcs_with_0_or_2_mg/6-xanthine/3-55NaCl_Mg/1-40winCmplx_30winLig/1-rep1/1-sys_prep’`
2. Run the script `‘0-add_Mg.tcl’` using:
   `‘vmd -dispdev text -e 0-add_Mg.tcl’`
   This script places $Mg^{2+}_{I}$ and $Mg^{2+}_{II}$ at the center of mass of the coordinating oxygens of the backbone RNA. For a list of the coordinating oxygens, see section 2.1. The outputs are `‘mg1.pdb’`, `‘mg2.pdb’`, and `‘mg3.pdb’`, which specify $Mg^{2+}$ coordinates. Only `‘mg1.pdb’` ($Mg^{2+}_{I}$) and `‘mg3.pdb’` ($Mg^{2+}_{II}$) eventualy get placed in the system for calculations with two $Mg^{2+}$ ions as specified in `‘tleap.in’`.
3. Run the script `‘1-ligand_fit.tcl’` using:
   `‘vmd -dispdev text -e 1-ligand_fit.tcl’`
   This script outputs the RNA coordinates into `‘rna.pdb’`, then after aligning the xanthine to the bound theophylline, outputs aligned xanthine coordinates to `‘lig.pdb’`.
4. Create a conda environment and install ambertools:
   `‘conda create --name ambertools’`
   `‘conda activate ambertools’`
   `‘conda install -c conda-forge ambertools’`

5. Run the script `‘2-run_system_setup.sh’` using:
   `‘./2-run_system_setup.sh’`
   In this script, first the ligand’s charge is assigned and then antechamber is deployed to generate ligand’s force field parameters. Then the `‘tleap.in’` script is executed, which loads in the force field parameters as well as the RNA, xanthine, and $Mg^{2+}$ coordinates. After adding a water box with 55 mM NaCl (9 $Na^{+}$ and $Cl^{+}$ ions, system’s coordinate (`‘box.pdb’`) and topology (`‘box.prmtop’`) files are written out. These two files can be directly used in NAMD.

#### 8.1.2 Pre-BFEE2 equilibration

**Aim**: Equilibrate the ligand-bound system before setting up the free energy calculations. We use the last frame from this equilibration as a start point in the BFEE2 alchemical free energy calculation procedure.

1. Go to the system simulation directory:
   `‘cd ./fe_calcs_with_0_or_2_mg/6-xanthine/3-55NaCl_Mg/1-40winCmplx_30winLig/1-rep1/2-sim_run’`
2. Set up the restraint reference files:
   `‘cd ./restraints’`
   `‘vmd -dispdev text -e restraints.tcl’`
   `‘restraints.tcl’` creates a reference pdb `‘restraints.pdb’` in which all the heavy atoms of RNA, ligand, and

the $Mg^{2+}$ ions are marked by setting their beta column to 1. Every other atom in the reference file has beta value of 0.

3. Run the ‘`run_1.sh`’ script found in ‘`2-sim_run`’ to start the pre-BFEE2 equilibration simulations to be executed sequentially.
   Please note that in ‘`run_1.sh`’, the hardcoded NAMD3 path (or ‘`$NAMD3_PATH`’) should be replaced with the path to the NAMD3 binary file that can be download from:
   ‘`https://www.ks.uiuc.edu/Development/Download/download.cgi?PackageName=NAMD`’
   Depending on the computing resources available one needs to use a compatible NAMD build.
   In this study we used p3.2xlarge instances on AWS for pre-BFEE2 equilibration.
4. When the last step of the simulations is finished, then use the last frame to re-wrap the simulation box and center the RNA using:
   ‘`vmd -dispdev text -e wrap.tcl`’
   This script outputs the equilibrated system with the RNA centered in ‘`./ini/eq.pdb`’.

#### 8.1.3 BFEE2 setup and calculation

**Aim**: Use the last frame of the equilibrated system to setup alchemical free energy calculations using BFEE2.

1. Install BFEE2 in a new conda environment:
   ‘`conda create --name bfee`’
   ‘`conda activate bfee`’
   ‘`conda install -c conda-forge BFEE2`’

2. Open the BFEE2 GUI:
   ‘`cd ./fe_calcs_with_0_or_2_mg/6-xanthine/3-55NaCl_Mg/1-40winCmplx_30winLig/1-rep1/`
   ‘`BFEE2Gui.py`’

3. Enter the path to the topology file and the equilibrated pdb, choose the alchemical route and specify the number of windows for the ligand-bound and ligand only systems. See Fig. S2a, b for parameters selected through BFEE2 GUI.
4. Press ‘Generate Inputs’ and when prompted to select a directory location for writing, choose ‘`./2-sim_run`’.
5. Now we have a ‘BFEE’ directory in which all the input files for the alchemical free energy calculations are stored. We need to first neutralize the ligand only system, as this system is automatically generated with BFEE2 by removing the RNA and only keeping the ligand, water, and ions. Since RNA has negative charge, upon its removal, we need to re-neutralize the system.
   Activate the ambertools conda environment and use the ‘`tleap_ligOnly.in`’:
   ‘`cd BFEE`’
   ‘`conda activate ambertools`’
   ‘`tleap -f tleap_ligOnly.in >> tleap_ligOnly.out 2>&1`’
6. Since the ‘ligandOnly.pdb’ is changed in the previous step, we need to regenerated the reference, xyz, and index files for the ligand only system. Use ‘`gen_xyz_ligOnly.py`’ with bfee conda environment:
   ‘`conda activate bfee`’
   ‘`python gen_xyz_ligOnly.py`’

7. To start the simulations, we used the script ‘`run_1.sh`’ and ‘`run_2.sh`’. ‘`run_1.sh`’ starts all the independent backward simulations in parallel and ‘`run_2.sh`’ starts the forward simulations which are dependent on the output of backward simulations. You can find this script in ‘`BFEE`’ directory in the repository.
   ‘`./run_1.sh`’
   ‘`./run_2.sh`’
   To perform the BFEE2 simulations for each replica, we used the AWS g5.8xlarge instances and distributed the CPU cores between the independent runs, as can be found in ‘`run.sh`’.
   Depending on the computing resources available, and the number of windows for each step, one should change the number of CPU cores allocated to each simulation for more efficiency.

##### 8.1.4 Analysis

In order to do the post-processing of the BFEE2 simulations for multiple ligands, replicates, and conditions use the following script:

```
‘cd ./fe_calcs_with_0_or_2_mg/’
‘conda activate bfee’
‘python post_treatment.py’
```

Before running the script, please remember to adjust the hard coded main directory path (mainDir) to point at the location of ‘/fe_calcs_with_0_or_2_mg/’ directory as shown in the script. To analyze results of multiple ligands, replicates, and conditions, do not forget to adjust compound list (‘cmpnd_list’), condition list (‘cond_list’), and replicate list (‘rep_list’). You can edit the name of the output file to be informative for the selection you made. Analysis results are written to ‘./fe_calcs_with_0_or_2_mg/results/BFE_with_failed/’ as text files.

#### 8.2 Simulation protocol for systems with OPC water model

In this section, we describe the protocol for preparing and simulating systems with OPC water model as opposed to TIP3P, which is described in 8.1.

##### 8.2.1 System preparation with OPC water model

The files for the OPC system can be found in:
‘./fe_calcs_with_0_or_2_mg/6-xanthine/3-55NaCl_Mg/6-opc_40winCmplx_30winLig/1-rep1’.
To prepare the OPC system follow the same steps as section 8.1.1, as all the scripts for system generation are identical. The only exception is the ‘tleap.in’ script in which the OPC water model is used to solvate the system. Hence, please make sure to use the ‘tleap.in’ script found here:
‘./fe_calcs_with_0_or_2_mg/6-xanthine/3-55NaCl_Mg/6-opc_40winCmplx_30winLig/1-rep1/1-sys_prep/tleap.in’.
for the OPC systems.

##### 8.2.2 Pre-BFEE2 equilibration with OPC water model

Since NAMD3 does not support 4-site water models yet, we used CUDA accelerated NAMD2 for the Pre-BFEE2 equilibration of the OPC systems.

Use ‘./fe_calcs_with_0_or_2_mg/6-xanthine/3-55NaCl_Mg/6-opc_40winCmplx_30winLig/1-rep1/2-sim_run/run.sh’ and change the hard coded NAMD2 CUDA PATH (or ‘$NAMD2_CUDA_PATH’) with the CUDA accelerated NAMD2 binary compatible with your computing resources.
Please also note that the NAMD input configuration files for the OPC water model include the following line to use the 4-site water model:
‘watermodel tip4’

##### 8.2.3 BFEE2 setup and calculation with OPC water model

Follow the steps 1-4 of the section 8.1.3 for BFEE2 set up.
In step 5, use the tleap script ( ‘tleap_ligOnly_opc.in’) compatible with the OPC water model found in:
‘./fe_calcs_with_0_or_2_mg/6-xanthine/3-55NaCl_Mg/6-opc_40winCmplx_30winLig/1-rep1/2-sim_run/BFEE/’
Follow step 6 form the section 8.1.3.
Before running the BFEE2 simulations, one needs to add the line ‘watermodel tip4’ to all the NAMD configuration files generated by BFEE2. You can do this using:

```
‘cd ./fe_calcs_with_0_or_2_mg/6-xanthine/3-55NaCl_Mg/6-opc_40winCmplx_30winLig/1-rep1/2-sim_run/BFEE/’
‘sed -i ’3iwatermodel tip4’ */*.conf’
```

Now the input files are ready, and you can run calculations using ‘run_1.sh’ and ‘run_2.sh’ files sequentially. These files are found in:
‘./fe_calcs_with_0_or_2_mg/6-xanthine/3-55NaCl_Mg/6-opc_40winCmplx_30winLig/1-rep1/2-sim_run/BFEE/’.
Please note for the TI simulations, one can use the CUDA accelerated NAMD2. However, for the FEP steps, the non-CUDA accelerated NAMD2 needs to be used, which makes those calculations much slower.

##### 8.2.4 Analysis

Follow the same steps as in section 8.5.4.

#### 8.3 Simulation protocol for systems with OpenFF

In this section, we describe the protocol for preparing and simulating systems using OpenFF 2.0.0 as the force field describing interactions of the ligand, as opposed to GAFF2, which is described in 8.1.
**main directory**: '`./fe_calcs_with_3_mg/6-xanthine/1-55NaCl_3Mg/2-OpenFF_40winCmplx_30winLig/1-rep1/`'

##### 8.3.1 System preparation with OpenFF

Do all the steps (1-5) described in section 8.1.1, and generate system's coordinate and topology files and rename them to '`box_gaff.inpcrd`' and '`box_gaff.prmtop`', respectively. Next we use the generated coordinate and topology files and replace ligand's GAFF2 parameter with OpenFF 2.0.0, following the swap amber parameters example from the OpenFF-toolkit.
Alternative URL for finding the tutorial for replacing ligand parameters in an already-parametrized system can be found here: "Replacing ligand parameters in an already-parametrized system".
Navigate to:
'`cd ./fe_calcs_with_3_mg/6-xanthine/1-55NaCl_3Mg/2-OpenFF_40winCmplx_30winLig/1-rep1/1-sys_prep`'
Then you can use the '`gen_openFF.py`' script which uses the ParmEd package. Run:
'`python gen_openFF.py`'
and outputs '`box.inpcrd`' and '`box.prmtop`', with the OpenFF force field for the ligand.
Run the '`fix_openff_pdb.tcl`' script to rewrite PDB file with VMD:
'`vmd -dispdev text -e  fix_openff_pdb.tcl`'
This step fixes any atom indexing issues introduced by '`gen_openFF.py`' script earlier.

##### 8.3.2 Pre-BFEE2 equilibration with OpenFF

Use the same steps as section 8.1.2.

##### 8.3.3 BFEE2 setup and calculation with OpenFF

Follow the same steps as section 8.1.3.

##### 8.3.4 Analysis

Follow the same steps as in section 8.5.4.

#### 8.4 Simulation protocol for systems with RNA backbone restraints

In this section, we describe the protocol for preparing and simulating systems with RMSD backbone restraints on the RNA.
**main directory**: '`./fe_calcs_with_0_or_2_mg/6-xanthine/8-5NaCl_Mg_bb_colvar/1-40winCmplx_30winLig/1-rep1`'.

##### 8.4.1 System preparation with RNA backbone restraints

Use the same steps as section 8.1.1.

##### 8.4.2 Pre-BFEE2 equilibration with RNA backbone restraints

1. Go to the system simulation directory:
   '`cd ./fe_calcs_with_0_or_2_mg/6-xanthine/8-5NaCl_Mg_bb_colvar/1-40winCmplx_30winLig/1-rep1/2-sim_run`'
2. Set up the restraint reference file:
   '`cd ./restraints`'
   '`vmd -dispdev text -e restraints.tcl`'
   '`restraints.tcl`' creates a reference pdb '`restraints.pdb`' in which all the heavy atoms of RNA, ligand, and the $Mg^{2+}$ ions are marked by setting their beta column to 1. Every other atom in the reference file has beta value of 0.

3. The RMSD backbone restraint is applied using the COLVAR module in NAMD. Set up the necessary files using:
   ‘vmd -dispdev text -e gen_index_group.tcl’
   ‘python gen_xyz_bb_colvar.py’

4. Run the ‘run_1.sh’ script found in ‘2-sim_run’ to start the pre-BFEE2 equilibration simulations to be executed sequentially.

#### 8.4.3 BFEE2 setup and calculation with RNA backbone restraints

Follow the steps 1-6 similar to section 8.1.3. Next we need to add RMSD backbone restraints to the FEP and TI calculations of the ligand-RNA complex. For this we first need to append the content of ‘restraints/bb.ndx’ file to the ned of ‘complex.ndx’. Second step is, modifying ‘colvars.in’ files of each BFEE step: insert ‘RMSD_bb’ colvar block as shown in ‘./fe_calcs_with_0_or_2_mg/automation_scripts/run_bb_colvar_BFEE.sh‘. Please remember to adjust ligand list (dir_list), condition list (cond_list), and replicate list (rep_list) before running this automation script. It also needs to be moved to one directory up (‘./fe_calcs_with_0_or_2_mg/‘) before running. As as result the following colvars.in files must are modified:

1. 000_eq/colvars.in.
2. 001_MoleculeBound/colvars.in
3. 002_RestraintBound/colvars_backward.in
4. 002_RestraintBound/colvars_forward.in

Then BFEE calculations are ready to be run with: ‘./run_1.sh’

#### 8.4.4 Contribution of the RNA backbone restraints

When RNA backbone restraints are applied, we need to perform an extra step, corresponding to step 5 of the thermodynamic cycle in Fig. 2d. In this extra step, we first apply RMSD backbone restraints on the RNA only system and allow it to equilibrate for 100 ns. Next, we use TI to reversibly turn off and turn on the restraints and calculate their contribution.
**main directory**: ‘./fe_calcs_with_0_or_2_mg/8-rna_RMSD_colvar_contr/1-55NaCl_2Mg/1-40win/1-rep1’.

Follow these steps to set up the input files:

1. ‘cd ./fe_calcs_with_0_or_2_mg/8-rna_RMSD_colvar_contr/1-55NaCl_2Mg/1-40win/1-rep1/1-sys_prep’

2. ‘vmd -dispdev text -e 0-add_Mg.tcl’

3. ‘vmd -dispdev text -e 1-extract_lig_pdb_resname.tcl’

4. Create a conda environment and install ambertools:
   ‘conda create --name ambertools’
   ‘conda activate ambertools’
   ‘conda install -c conda-forge ambertools’

5. Run system setup:
   ‘./2-run_system_setup.sh’

Next, setup restraints and run the equilibrium simulations:

1. ‘cd ./fe_calcs_with_0_or_2_mg/8-rna_RMSD_colvar_contr/1-55NaCl_2Mg/1-40win/1-rep1/2-sim_run’

2. ‘cd ./restrasints’ ‘vmd -dispdev text -e restraints.tcl’
   ‘vmd -dispdev text -e gen_index_group.tcl’
   ‘python gen_xyz_bb_colvar.py’

3. ‘cd ../’
4. ‘./run_1.sh’
5. After the equilibration is finished:
   ‘vmd -dispdev text -e wrap_rna_only.tcl’

Next, setup and run the TI simulation:

1. ‘cd ./fe_calcs_with_0_or_2_mg/8-rna_RMSD_colvar_contr/1-55NaCl_2Mg/1-40win/1-rep1/2-sim_run/’

2. ‘./run_contr.sh’

TI output will be collected in ‘contribution_RMSD_ref_wrap/’ directory.

#### 8.4.5 Analysis

Follow the same steps as in section 8.5.4.
For the analysis of the simulations regarding the contribution of the RNA backbone restraints, use the following script: ‘python post_treatment_rnaOnly.py’ with a single change in ‘postTreatment.py’ script to accept eight collective variables, instead of the original seven.

### 8.5 Simulation protocol for doubling the sampling

Since the highest variation between replicas are from step 3 of the thermodynamic cycle in Fig. 2, we explored doubling the sampling for this step and check for possible improvements. Please note that we only re-simulated step 3 and to report a final binding free energy, we used the base calculation that was already performed for all the other steps.
**main directory 80 windows, 1 ns**: ‘./fe_calcs_with_0_or_2_mg/6-xanthine/3-55NaCl_Mg/2-80winCmplx/1-rep1’.
**main directory 40 windows, 2 ns**: ‘./fe_calcs_with_0_or_2_mg/6-xanthine/3-55NaCl_Mg/5-40winCmplx_2ns/1-rep1’.

#### 8.5.1 System preparation for doubling the sampling

For these calculations, we can use the same coordinate and topology files that we created in section 8.1.1. You can find the ‘box.pdb’ and ‘box.prmtop’ in the ‘1-sys_prep’ directory of both 80 windows and 40 windows of 2 ns.

#### 8.5.2 Pre-BFEE2 equilibration for doubling the sampling

This step is not performed since we only simulate step 3 of the thermodynamic cycle in Fig. 2.

#### 8.5.3 BFEE2 setup and calculation with double the sampling

Next, we need to copy over the step 3 of the thermodynamic cycle in Fig. 2, from the base calculation and change the input file to double the sampling. You can find the BFEE2 directory corresponding to step 3 in:
‘2-sim_run/BFEE/001_MoleculeBound’. Since we copied this directory from the base calculation (40 windows, 1 ns), to run 80 windows, 1 ns we need to run the following commands:

1. sed -i ’s/runFEP 1.0 0.0 -0.025 500000/runFEP 1.0 0.0 -0.0125 500000/’
   ./2-80winCmplx/2-sim_run/BFEE/001_MoleculeBound/*conf

2. sed -i ’s/runFEP 0.0 1.0 0.025 500000/runFEP 0.0 1.0 0.0125 500000/’
   ./2-80winCmplx/2-sim_run/BFEE/001_MoleculeBound/*conf

In the case of 40 windows of 2 ns we need to run:

1. `sed -i 's/alchEquilSteps 100000/alchEquilSteps 200000/' ./40winCmplx_2ns/2-sim_run/BFEE/001_MoleculeBound/*conf`

2. `sed -i 's/runFEP 1.0 0.0 -0.025 500000/runFEP 1.0 0.0 -0.025 1000000/' ./40winCmplx_2ns/2-sim_run/BFEE/001_MoleculeBound/*conf`

3. `sed -i 's/runFEP 0.0 1.0 0.025 500000/runFEP 0.0 1.0 0.025 1000000/' ./40winCmplx_2ns/2-sim_run/BFEE/001_MoleculeBound/*conf`

Now you can run the simulations using:
'`cd ./BFEE/001_MoleculeBound/run.sh`'

#### 8.5.4 Analysis

In order to do the post-processing of the BFEE2 simulations, use the following script:
'`conda activate bfee`'
'`python post_treatment_doubling_sampling.py`'

### 8.6 Protocol for MM-GBSA Calculations

MM-GBSA calculations take advantage of 100 ns equilibiration simulations run prior to each free energy calculation with BFEE2. MM-GBSA calculation scripts are organized under '`mmgbsa_calc_with_0_or_2_mg/`' directory.

1. Set up conda environments for LOOS and Ambertools23, named LOOS and Ambertools23 respectively.
2. Copy the following script and input files to the path of the equilibrium trajectory of each ligand:

- '`run_mmgbsa.sh`'
- '`strip_cpptraj_dry.in`'
- '`strip_cpptraj_lig.in`'
- '`strip_cpptraj_rna.in`'
- '`loos_to_vmddcd.tcl`'
- '`mmgbsa.in`'
- '`box.prmtop`'

2. Run '`run_mmgbsa.sh`' one by one for all three replicates of each ligand: '`source run_mmgbsa.sh`'

4. Extract estimates of binding free energy from the '`FINAL_RESULTS_MMPBSA.dat`' output file.

### 8.7 Overlap assessment of ΔU probability distribution function

One of the main challenges of free energy methods, such as FEP, is assessing the convergence of free energy estimates [13]. Considering a reference state, A, and a target state, B, the FEP identity can be written as:

$$\exp(-\beta\Delta G) = <\exp(-\beta\Delta U)>_A \tag{5}$$

where $\beta = 1/k_B T$, with $k_B$ being the Boltzmann constant and T representing temperature [14]. $\Delta G$ represents the free energy difference associated with the process of going from state A to B, and $\Delta U$ is the difference in the potential energy of those states. The $<>_A$ denotes that the ensemble average is calculated over the microstates generated using state A ensemble. On the other hand, we can choose to simulate the target state, B, and perform the ensemble averaging over the microstates representing B. The FEP identity in this case can be written as:

$$\exp(\beta \Delta G) =< \exp(\beta \Delta U) >_B \tag{6}$$

Now, considering a bidirectional transformation, we can use equations 5 and 6 to get two estimates for free energy. The natural question to answer is which estimate to report or how to combine them to have a better free energy estimate. The Bennett Acceptance Ratio (BAR) method provides an optimal solution to this problem by providing an estimation of the free energy, with minimum variance, using data from both simulated ensembles [16]:

$$\Delta G = ln \frac{< f(-\Delta U + C) >_B}{< f(\Delta U - C) >_A} + C \tag{7}$$

where $f(x) = 1/[1 + \exp(x)]$ is the Fermi function, $\Delta U = U_B - U_A$, and C is a constant. In order to calculate the ensemble averages in equation 7, one can store pair of $\Delta U = U_B - U_A$ histograms, $P_A(\Delta U)$ and $P_B(\Delta U)$, observed when sampling from the B and A ensembles, respectively. The values of the $\Delta U$ stored in these histograms are then used to calculate the ensemble averages of the two complementary Fermi functions. If the the histograms are separated by a large gap, such as the case shown in Figure S1, then no value of C would simultaneously result in non-zero values from the two complementary Fermi functions. This is a limitation of the BAR method, as equation 7 has been derived assuming a large-sample regime, which can be ensured when the ensemble averages $< f(-\Delta U + C) >_B$ and $< f(\Delta U - C) >_A$ are large compared to unity. As a result, to have the best estimates from the BAR method, one needs to ensure reasonable overlap between the $P_A(\Delta U)$ and $P_B(\Delta U)$.

A typical approach to assess convergence in each $\lambda$-window is to plot probability distribution functions of forward and backward transformations (Figure S7a) and visually inspect for overlap problems [64]. But with $\lambda$-windows, three replicates, and six ligands inspecting these plots visually became a cumbersome task. For quicker diagnosis of problematic $\lambda$-windows with insufficient overlap, we calculated Kullback–Leibler divergence ($D_{KL}$) to quantify the difference between potential energy distributions and made use of $D_{KL}$ vs $\lambda$-window bar plots for quick visualization (Figure S7b).

In this study, $D_{KL}$ is used as a proxy to characterize the degree of overlap between the two probability distribution functions of potential energy difference ($\Delta U$ distributions of forward and backwards transformations) in each $\lambda$-window. $D_{KL}$ for comparing the two distributions of $\Delta U$ is calculated as follows:

$$D_{KL}(P||Q) = \int_{-\infty}^{\infty} p(x) log(\frac{p(x)}{q(x)})\, dx \tag{8}$$

P and Q correspond to $P_{fwd}(\Delta U)$ and $P_{bwd}(\Delta U)$, respectively. More specifically, we used the "symmetrized $D_{KL}$" defined as:

$$D_{KL} = \frac{1}{2}[D_{KL}(P||Q) + D_{KL}(Q||P)] \tag{9}$$

Small $D_{KL}$ values indicate that the two histograms are closer to each other and as a result have a better degree of overlap. These results are shown in Figures S7-S11.

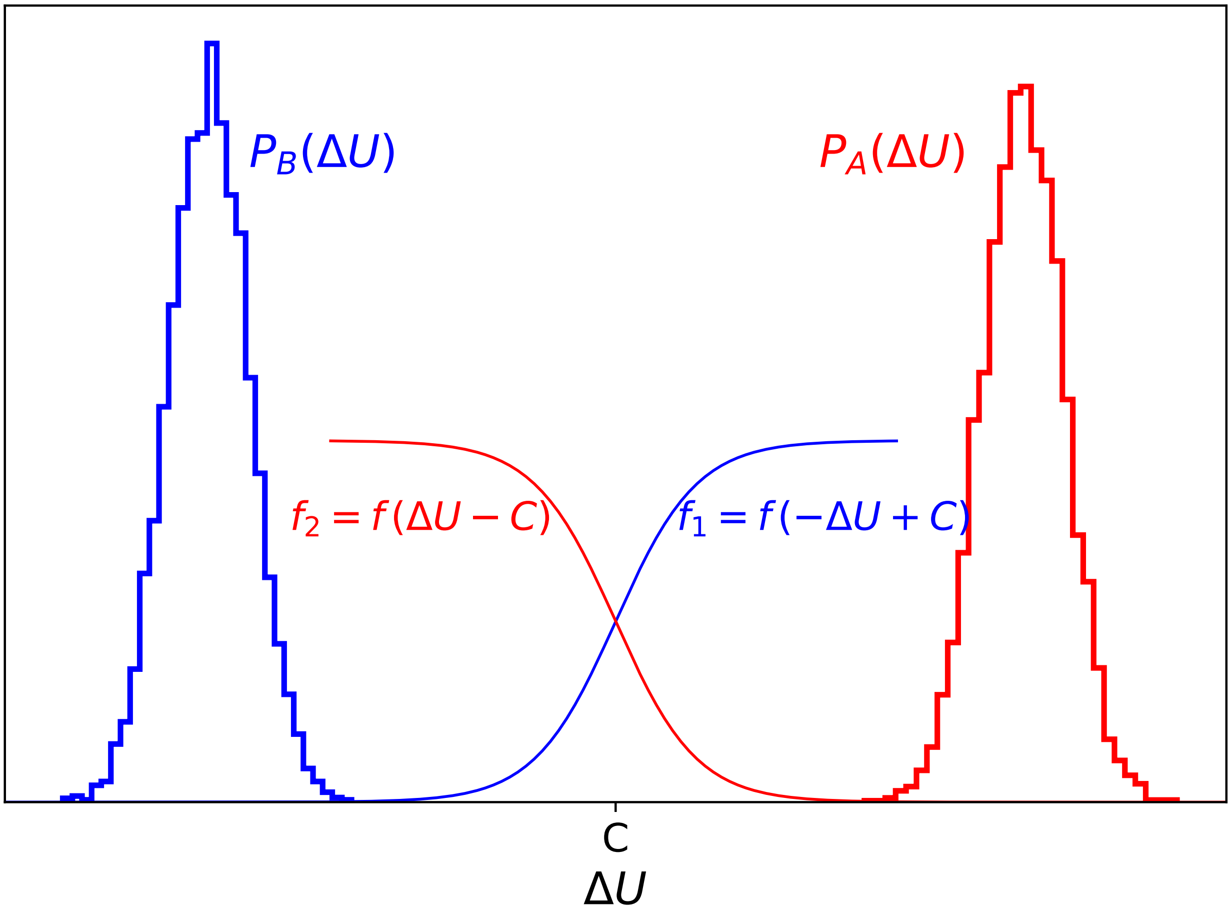


**Figure S1.** Histograms of $\Delta U$ values ($\Delta U = U_B - U_A$) sampled from the A and B ensembles, represented as $P_A(\Delta U)$ and $P_B(\Delta U)$, respectively. To determine the free energy difference between state A and B, according to equation 7, one must compute ensemble averages of complementary Fermi functions $f_1 = f(-\Delta U + C)$ and $f_2 = f(\Delta U - C)$. Due of a lack of overlap between $P_A(\Delta U)$ and $P_B(\Delta U)$, the widths of these Fermi functions are insufficient for achieving substantial overlap between $f_1$ with $P_B(\Delta U)$ and between $f_2$ and $P_A(\Delta U)$, simultaneously. This figure was adapted from Figure 5 of the article of Charles Bennett [16]. These are artificial histograms and do not represent actual data.

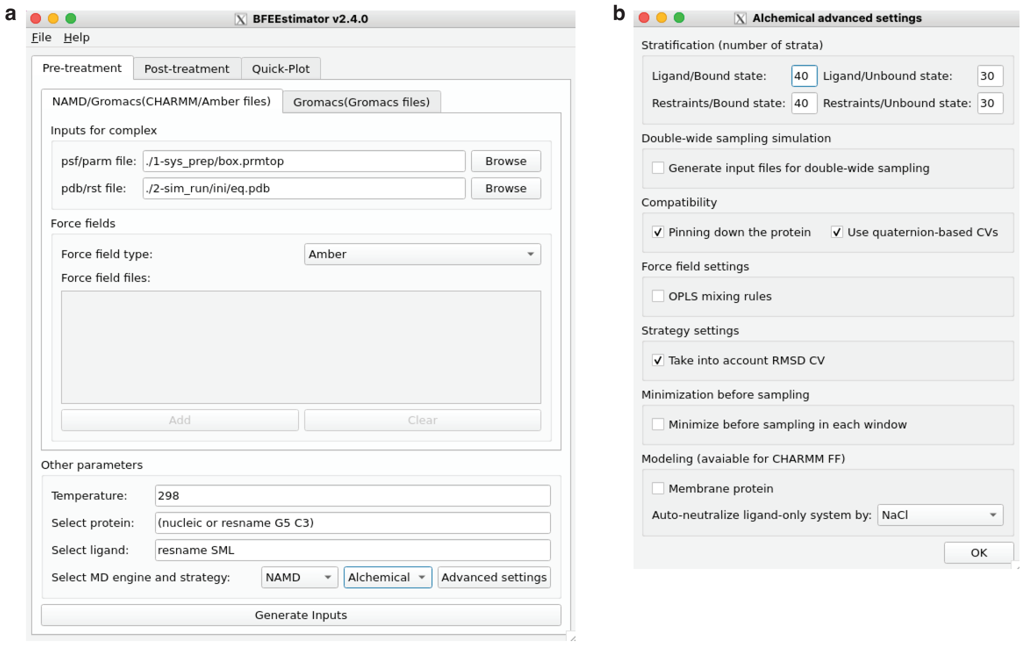


**Figure S2. Settings for the generation of inputs for the alchemical route using BFEE2 GUI . (a)** Enter the path to the topology file, and the equilibrated coordinates in the 'Inputs for complex' section. Select the RNA and ligand in the 'Select protein' and 'Select ligand', using the MDAnalysis syntax. **(b)** In the 'Advanced settings' set the number of window for the 'Bound state' and 'Unbound state'. Choose to 'Use quaternion-based CVs'

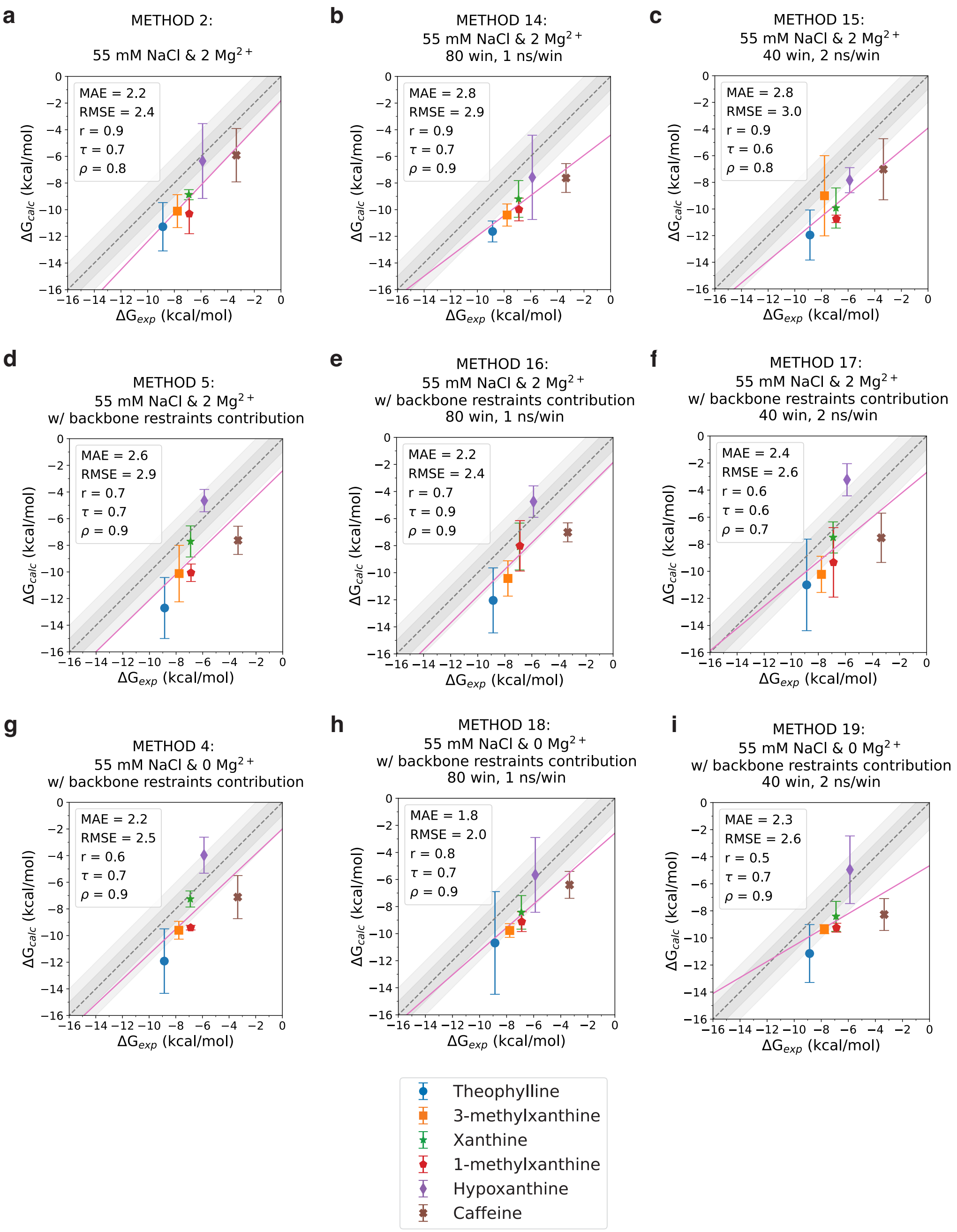


**Figure S3. Effects of doubling the sampling on binding free energy calculations** To check the convergence of the free energy calculations we double the sampling only in step 3 of the thermodynamic cycle Fig. 2b, and d. **(a)**, **(d)**, and **(g)** Base calculations, 40 windows and 1 ns/win, with 55 mM NaCl and 2 $Mg^{2+}$, 55 mM NaCl and 2 $Mg^{2+}$ and RNA backbone RMSD restraints, and 55 mM NaCl and 0 $Mg^{2+}$ and RNA backbone RMSD restraints. **(b)**, **(e)**, and **(h)** Doubling the sampling in step 3 by doubling the number of $\lambda$ windows to 80. **(c)**, **(f)**, and **(i)** Doubling the sampling in step 3 by doubling the sampling time per window to 2 ns/win. When doubling the sampling in step 3, all other steps are used from the base calculation.

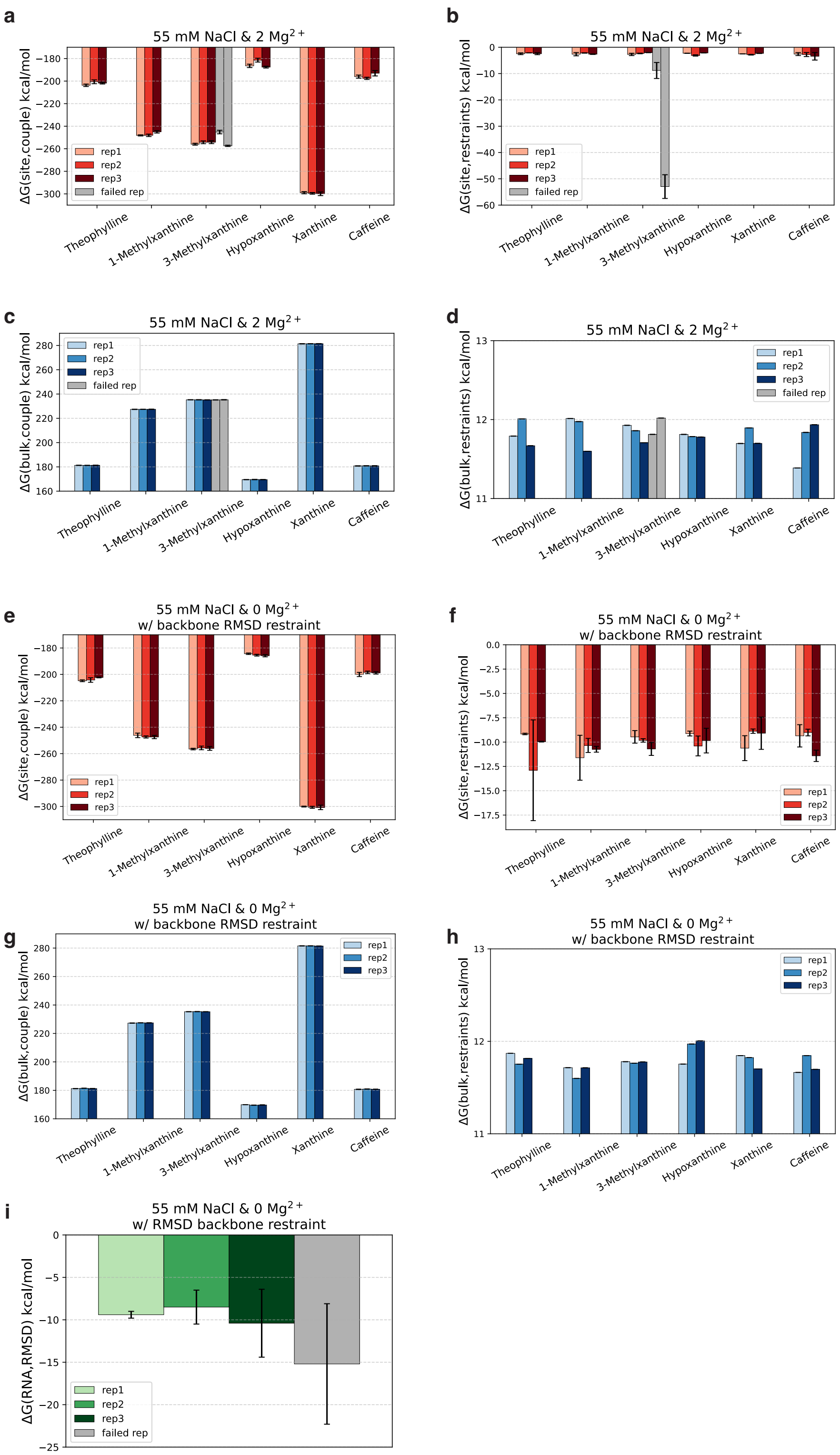


**Figure S4. Contributions of different BFEE2 steps in the absolute binding free energy. (a)**, **(b)**, **(c)**, **(d)** Free energy values for the system with 55 mM NaCl and 2 $Mg^{2+}$ (Method 2 in Table 1), related to steps 3, 4, 2, and 1 in Fig. 2b. **(e)**, **(f)**, **(g)**, **(h)** Free energy values for the system with 55 mM NaCl and 0 $Mg^{2+}$ with RMSD backbone restraints (Method 4 in Table 1), related to steps 3, 4, 2, and 1in Fig. 2d. **(i)** Step 5 of thermodynamic cycle: Contribution of backbone restraints on RNA-only system. Shades of red, blue, and green indicated accepted replicates. Grey color indicates rejected replicates.

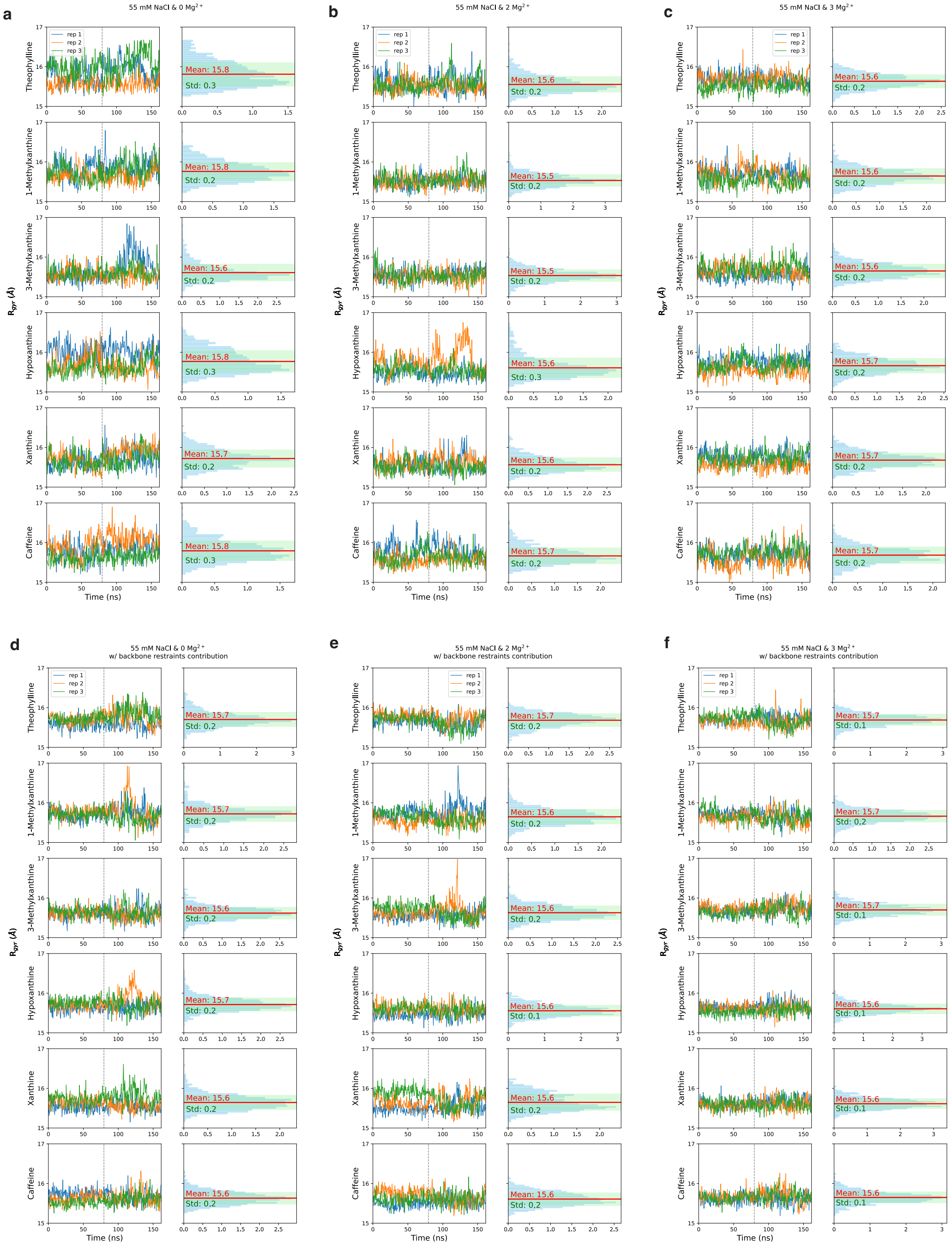


**Figure S5. RNA heavy atom radius of gyration ($R_{gyr}$ ). (a)** Time evolution of the radius of gyration of the RNA heavy atoms. Each independent replica of the system, under the condition of Method 1 (55 mM NaCl and 0 $Mg^{2+}$) , is represented by different colors in the left column. The right column displays a histogram, summarizing the aggregate data from three replicas, with mean and standard deviation values for the $R_{gyr}$ distributions in red and green, respectively. Each row corresponds to $R_{gyr}$ of RNA when bound to a different ligand. Subpanels **(b)** through **(f)** follow the same format as **(a)** but represent systems under different conditions: **(b)** Method 2 (55 mM NaCl and 2 $Mg^{2+}$ ), **(c)** Method 3 (55 mM NaCl and 3 $Mg^{2+}$ ), **(d)** Method 4 (55 mM NaCl and 0 $Mg^{2+}$ with backbone restraints contribution), **(e)** Method 5 (55 mM NaCl and 2 $Mg^{2+}$ with backbone restraints contribution), and **(f)** Method 6 (55 mM NaCl and 3 $Mg^{2+}$ with backbone restraints contribution).

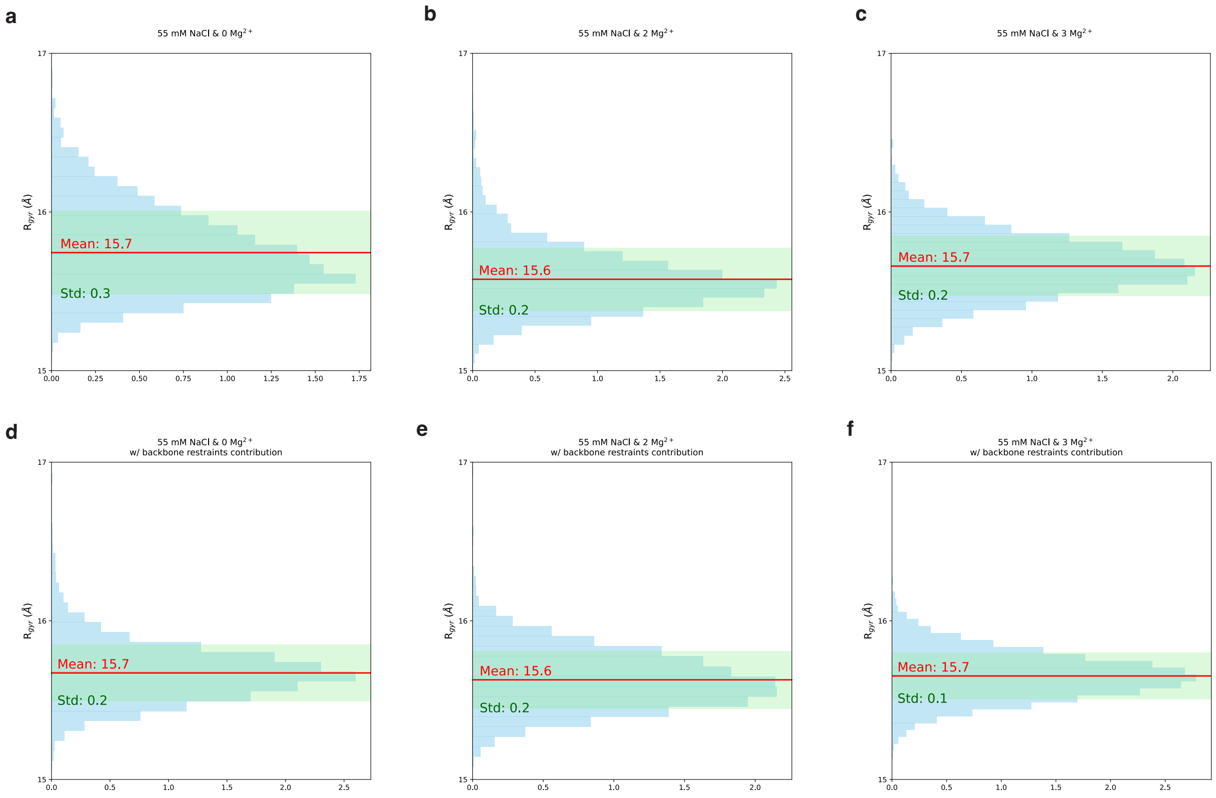


**Figure S6. Histograms of radius of gyration ($R_{gyr}$) calculated based on RNA heavy atoms.** Histograms depicting the distribution of the RNA heavy atom radius of gyration ($R_{gyr}$) across a range of system conditions, calculated from aggregate of three replicates for each system. Panels **(a)** through **(f)** correspond to different system conditions as follows: **(a)** Method 1 (55 mM NaCl and 0 $Mg^{2+}$ ), **(b)** Method 2 (55 mM NaCl and 2 $Mg^{2+}$ ), **(c)** Method 3 (55 mM NaCl and 3 $Mg^{2+}$ ), **(d)** Method 4 (55 mM NaCl and 0 $Mg^{2+}$ with backbone restraints contribution), **(e)** Method 5 (55 mM NaCl and 2 $Mg^{2+}$ with backbone restraints contribution), and **(f)** Method 6 (55 mM NaCl and 3 $Mg^{2+}$ with backbone restraints contribution). Each panel illustrates the aggregate $R_{gyr}$ distribution for the corresponding condition, considering all six ligands investigated in this study. The mean and standard deviation of the $R_{gyr}$ distribution for each condition are highlighted in red and green, respectively.

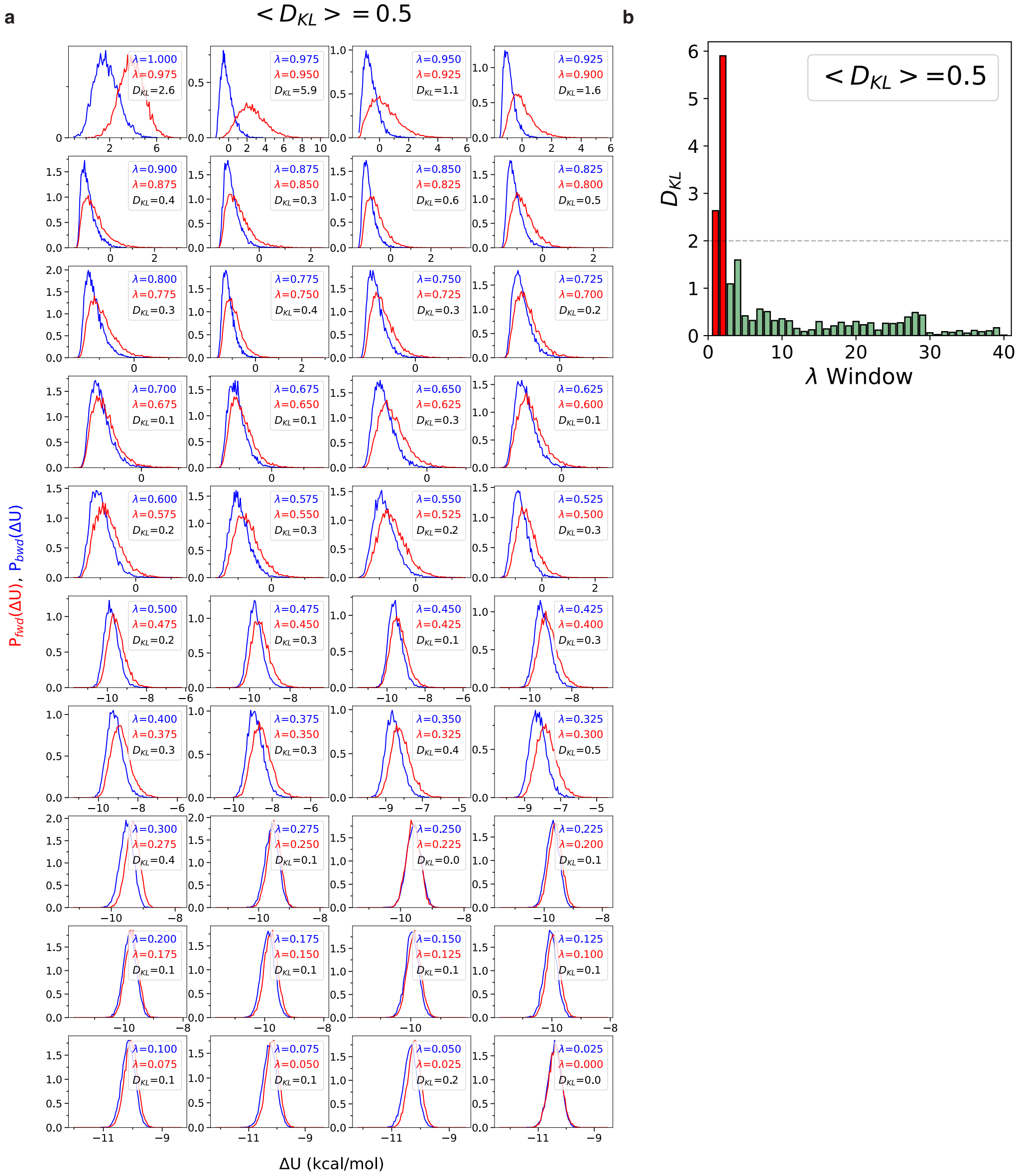


**Figure S7. Assessing overlap in probability distribution functions of potential-energy difference ( $\Delta U$). (a)** Examination of the extent of overlap between the probability distribution functions of potential-energy differences, $P(\Delta U)$, pertinent to the step 3 of Figure 2a. As step 3 is performed in a bidirectional manner with 40 steps in each direction, each subplot represents one of these 40 $\lambda$ windows, corresponding to two adjacent $\lambda$ values, shown in blue and red in the legend. In the backward transformation, the equilibrium ensemble is generated using the larger $\lambda$, and $\Delta U$ is calculated between the two $\lambda$ states. In the forward transformation the smaller $\lambda$ value generates the equilibrium ensemble. $P_{bwd}(\Delta U)$ represents the probability distribution of $\Delta U$ in the backward transformation (in blue), while $P_{fwd}(\Delta U)$ depicts the forward transformation (in red). To quantify the extent of overlap, we compute the symmetrized Kullback-Leibler (KL) divergence ($D_{KL}$) between the two probability distribution functions for each window. These $D_{KL}$ values are detailed in the subplot legends. The average $D_{KL}$ across all 40 windows, denoted as $< D_{KL} >$; on top of the plot, servers as a metric to gauge the transformation's overall quality and consistency. This panel shows the results for the first replica of the system with theophylline, 55 mM NaCl, and 0 $Mg^{2+}$ (Method 1). **b** Bar plot of the $D_{KL}$ values for all the 40 windows, with color-coding for clarity. $D_{KL}$ values exceeding 2 are portrayed in red, signifying regions of notable discrepancy. Conversely, values falling below or equal to 2 are represented in green, denoting regions with a higher degree of overlap, signifying enhanced reliability within the FEP analysis.

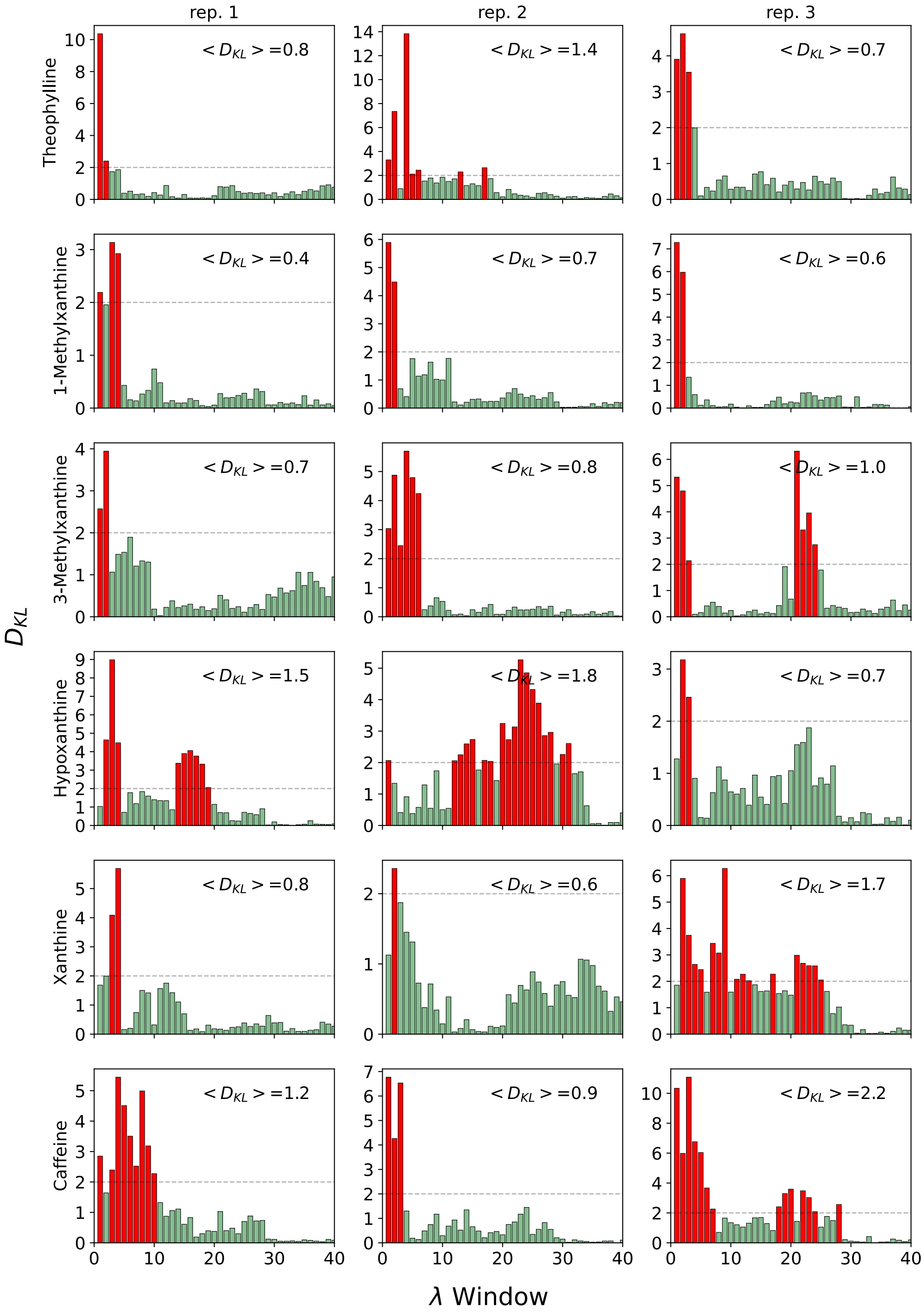


**Figure S8. Overlap analysis of $P_{bwd}(\Delta U)$ and $P_{fwd}(\Delta U)$ distributions with $D_{KL}$ for Method 2.** Similar to Figure S S6b, each subplot in this figure presents a bar plot illustrating the symmetrized Kullback-Leibler (KL) divergence ($D_{KL}$) between probability distribution of $\Delta U$ in the backward and forward transformations. All systems depicted here are simulated with 55 mM NaCl and 2 $Mg^{2+}$, without backbone restraints (Method 2). The transformation consists of 40 windows, each with 1 ns/window sampling time. Rows correspond to systems bound to various ligands, while columns show the three independent replicas for each system.

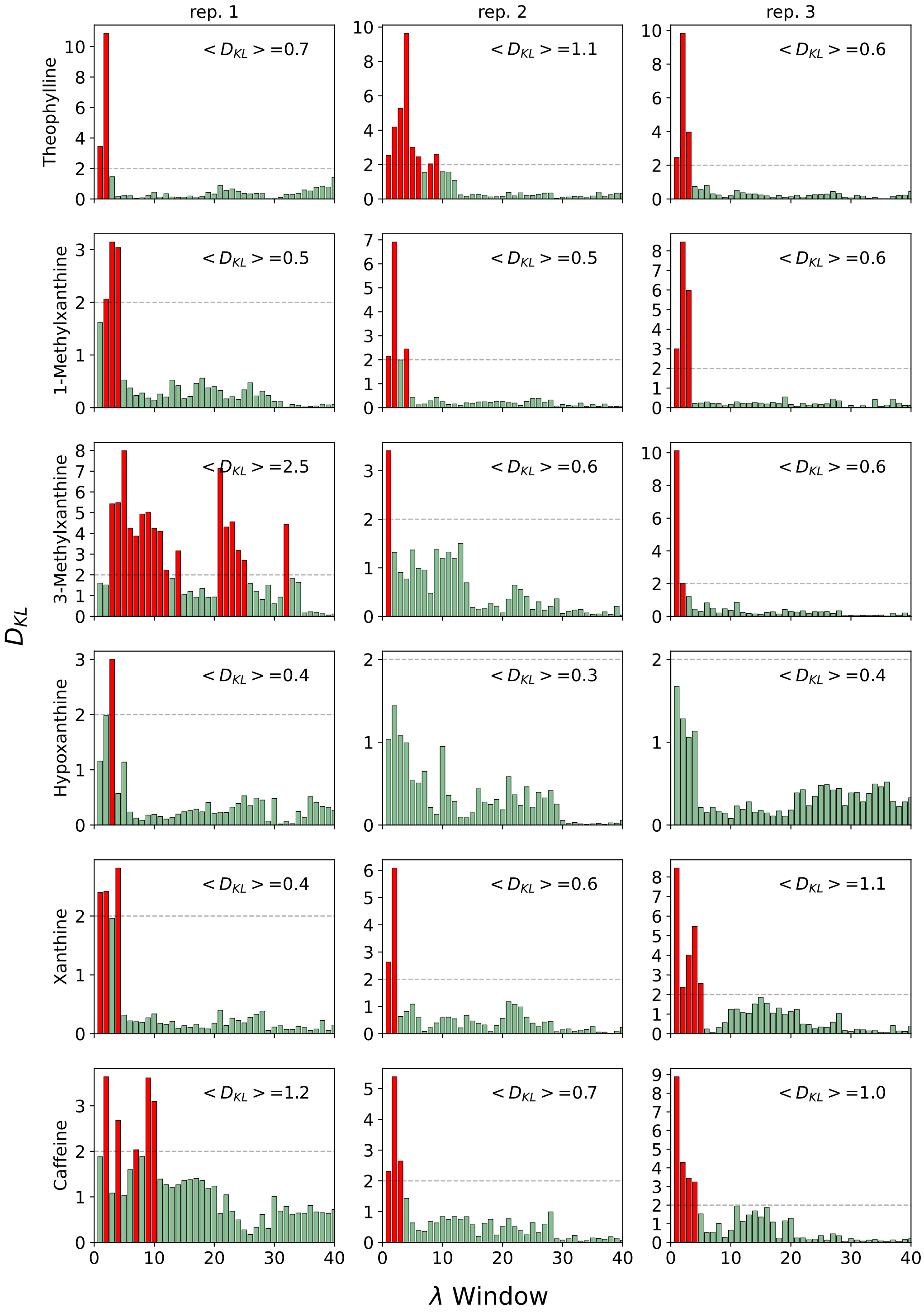


**Figure S9. Overlap analysis of $P_{bwd}(\Delta U)$ and $P_{fwd}(\Delta U)$ distributions with $D_{KL}$ for Method 15.** Similar to Figure S8, but for systems with 55 mM NaCl and 2 $Mg^{2+}$, without backbone restraints with 40 windows, each with 2 ns/window sampling time.

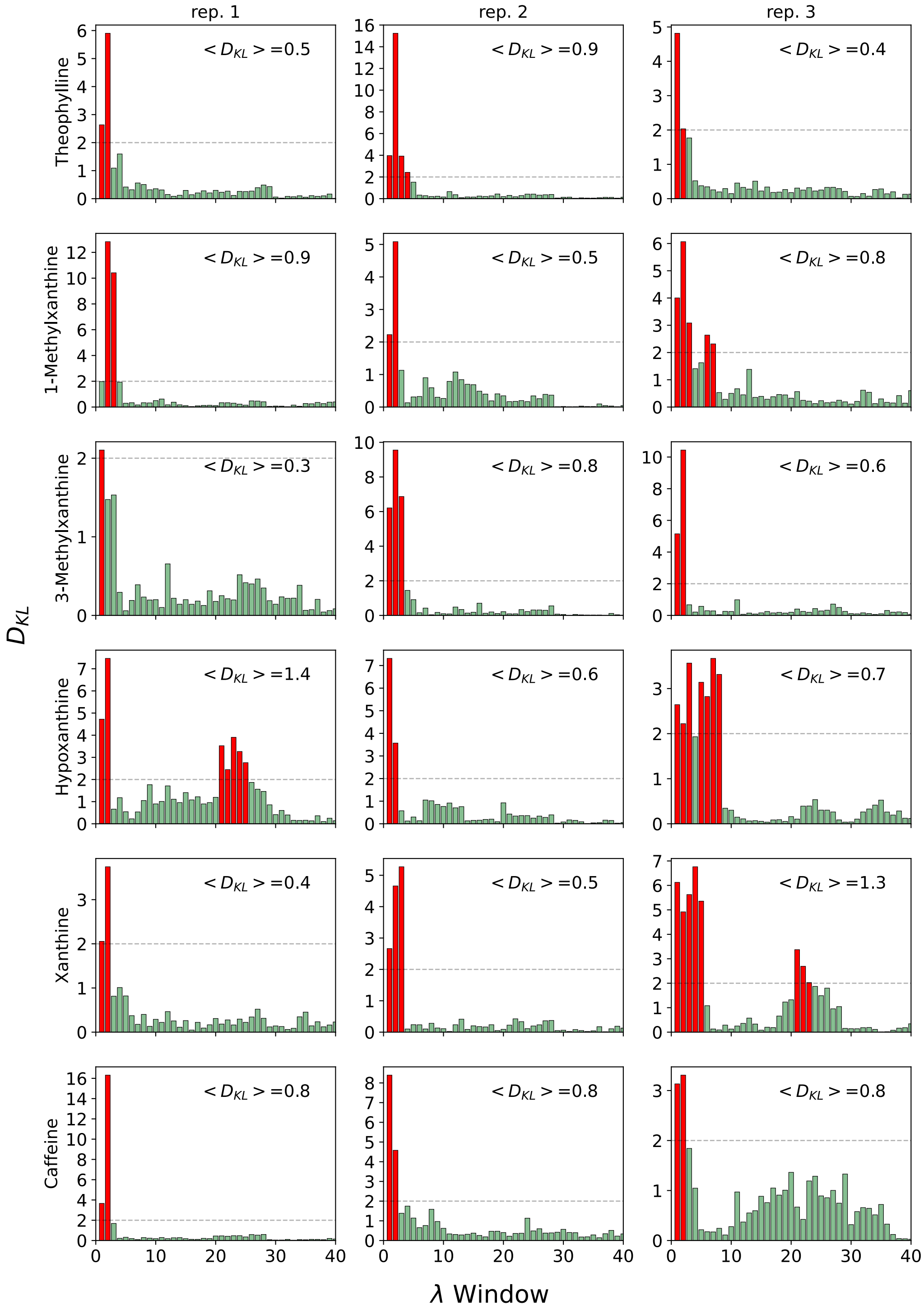


**Figure S10. Overlap analysis of $P_{bwd}(\Delta U)$ and $P_{fwd}(\Delta U)$ distributions with $D_{KL}$ for Method 4.** Similar to Figure S8, but for systems with 55 mM NaCl and 0 $Mg^{2+}$, with 40 windows, each with 1 ns/window sampling time and with backbone restraint contributions.

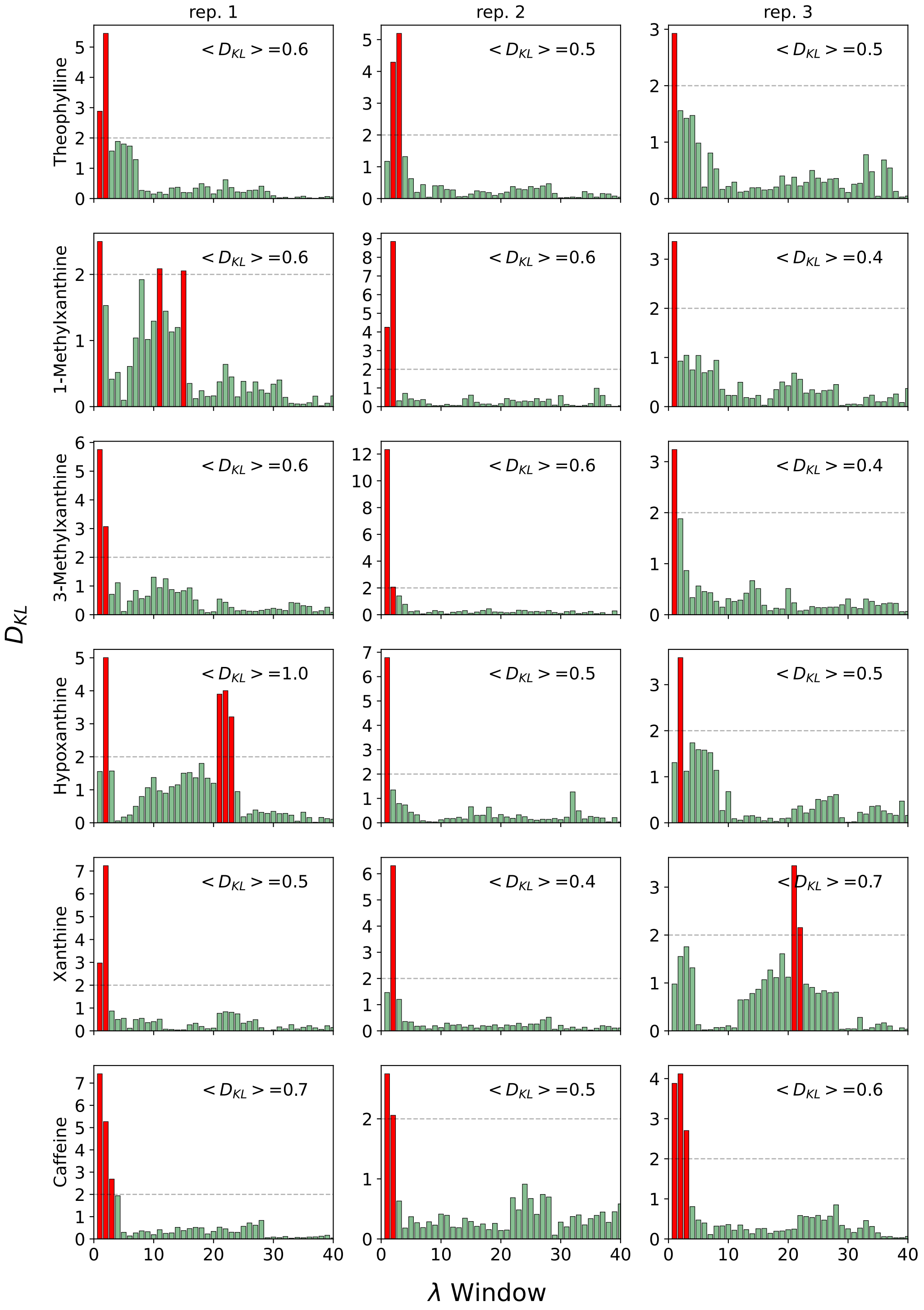


**Figure S11. Overlap analysis of $P_{bwd}(\Delta U)$ and $P_{fwd}(\Delta U)$ distributions with $D_{KL}$ for Method 19.** Similar to Figure S8, but for systems with 55 mM NaCl and 0 $Mg^{2+}$, with 40 windows, each with 2 ns/window sampling time and with backbone restraint contributions.

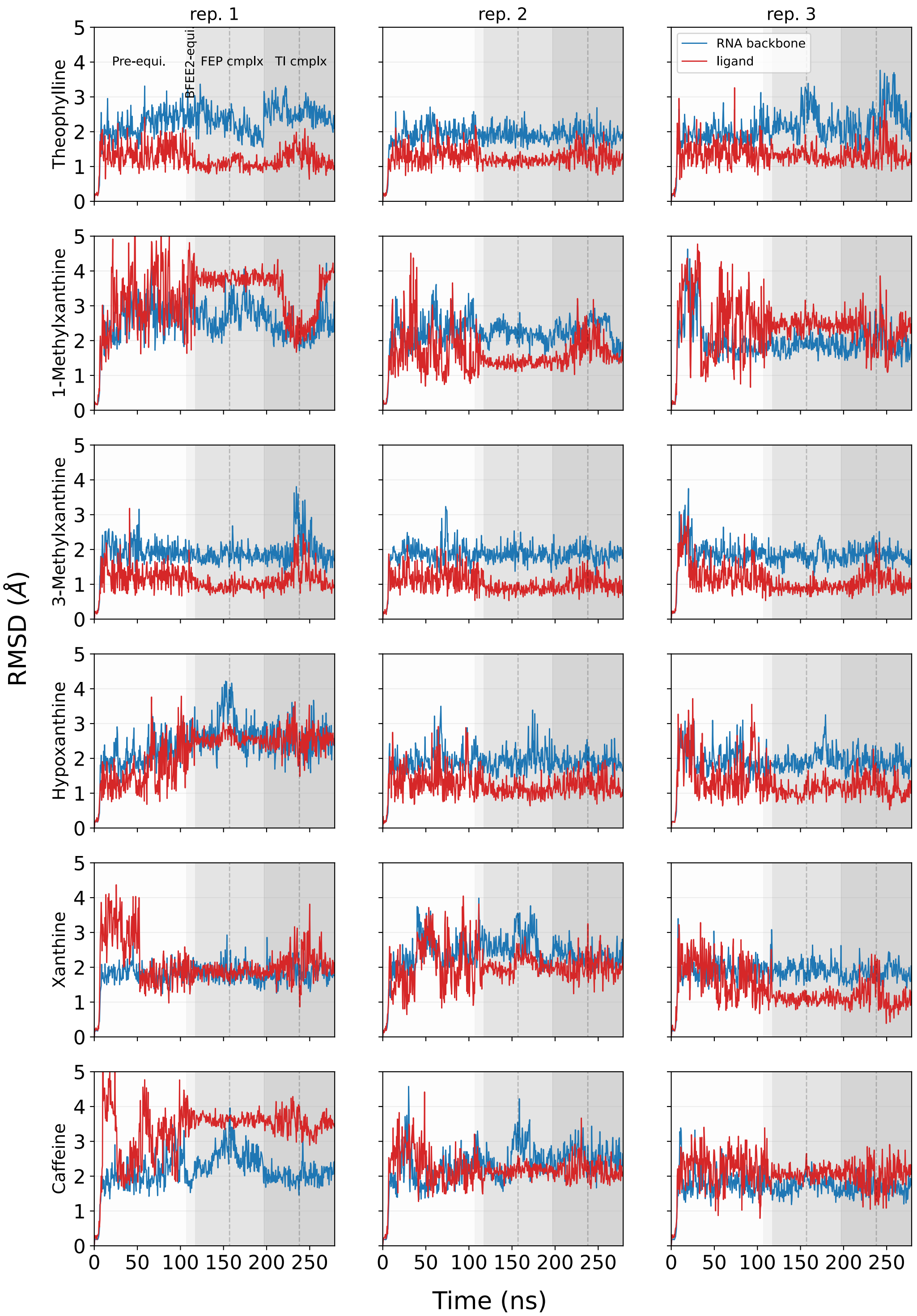


**Figure S12. Stability analysis of the RNA and the bound small molecule for Method 1.** Time evolution of the RMSD for RNA backbone heavy atoms (in blue) and the bound ligand (in red) under salt condition of 55 mM NaCl and 0 $Mg^{2+}$. Each subplot corresponds to a set of alchemical free energy calculations, with rows representing systems bound to various ligands, and columns signifying three independent replicas for each system. The background color of the plots is used to differentiate between different steps of the free energy calculation. The subplot on the top left, shows what each background color represents: "Pre-equi." represents equilibration steps prior to the BFEE2 protocol, "BFEE2-equi" depicts the 10 ns equilibration in the BFEE2 protocol, "FEP cmplx" corresponds to step 3 of Figure 2a, with a vertical dotted line distinguishing the backward and forward transformations, and "TI cmplx" corresponds to step 4 of Figure 2a, with another vertical dotted line distinguishing the backward and forward transformations.

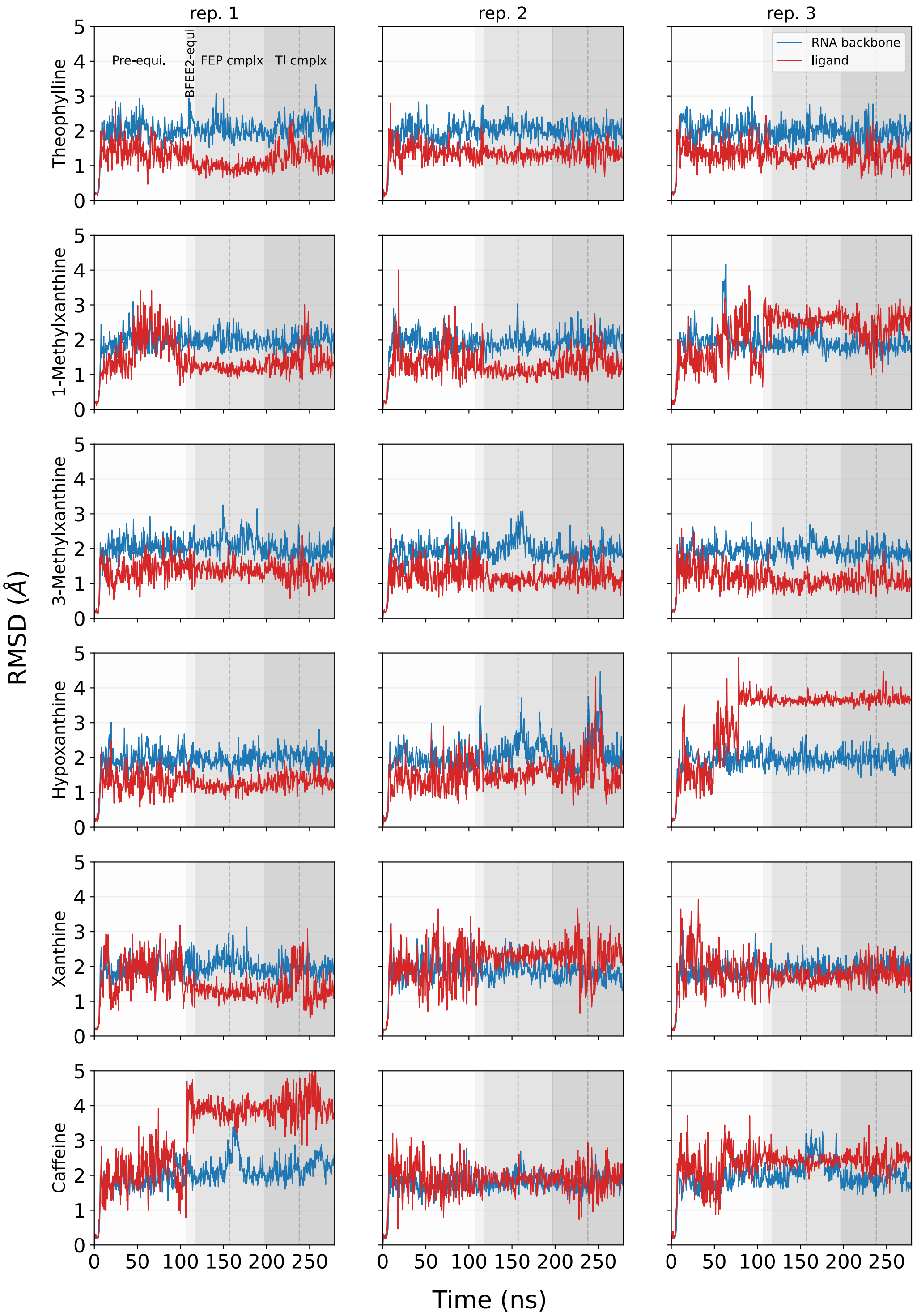


**Figure S13. Stability analysis of the RNA and the bound small molecule for Method 2** Similar to Figure S12 but for systems with 55 mM NaCl and 2 $Mg^{2+}$.

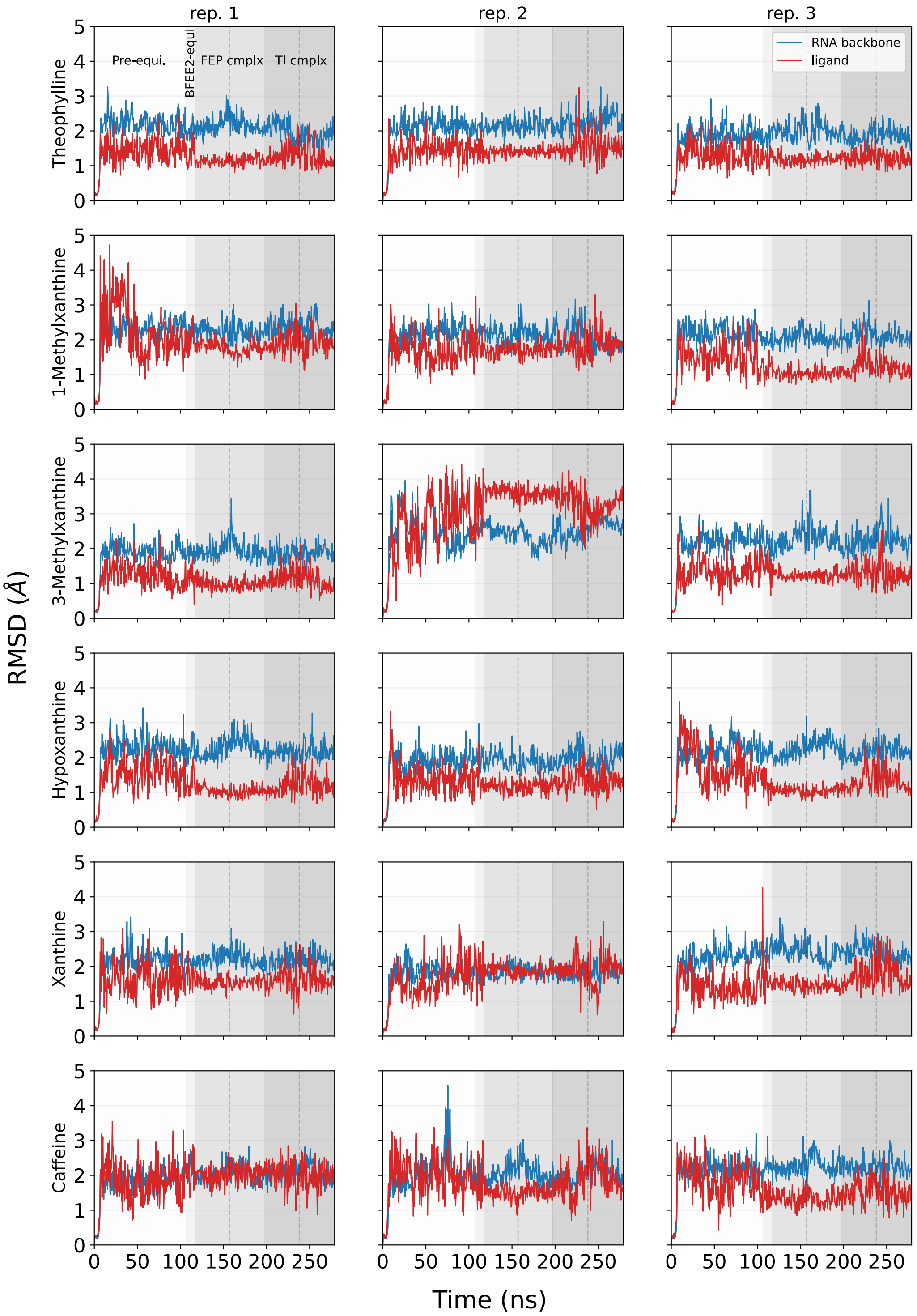


**Figure S14. Stability analysis of the RNA and the bound small molecule for Method 3.** Similar to Figure S12 but for systems with 55 mM NaCl and 3 $Mg^{2+}$.

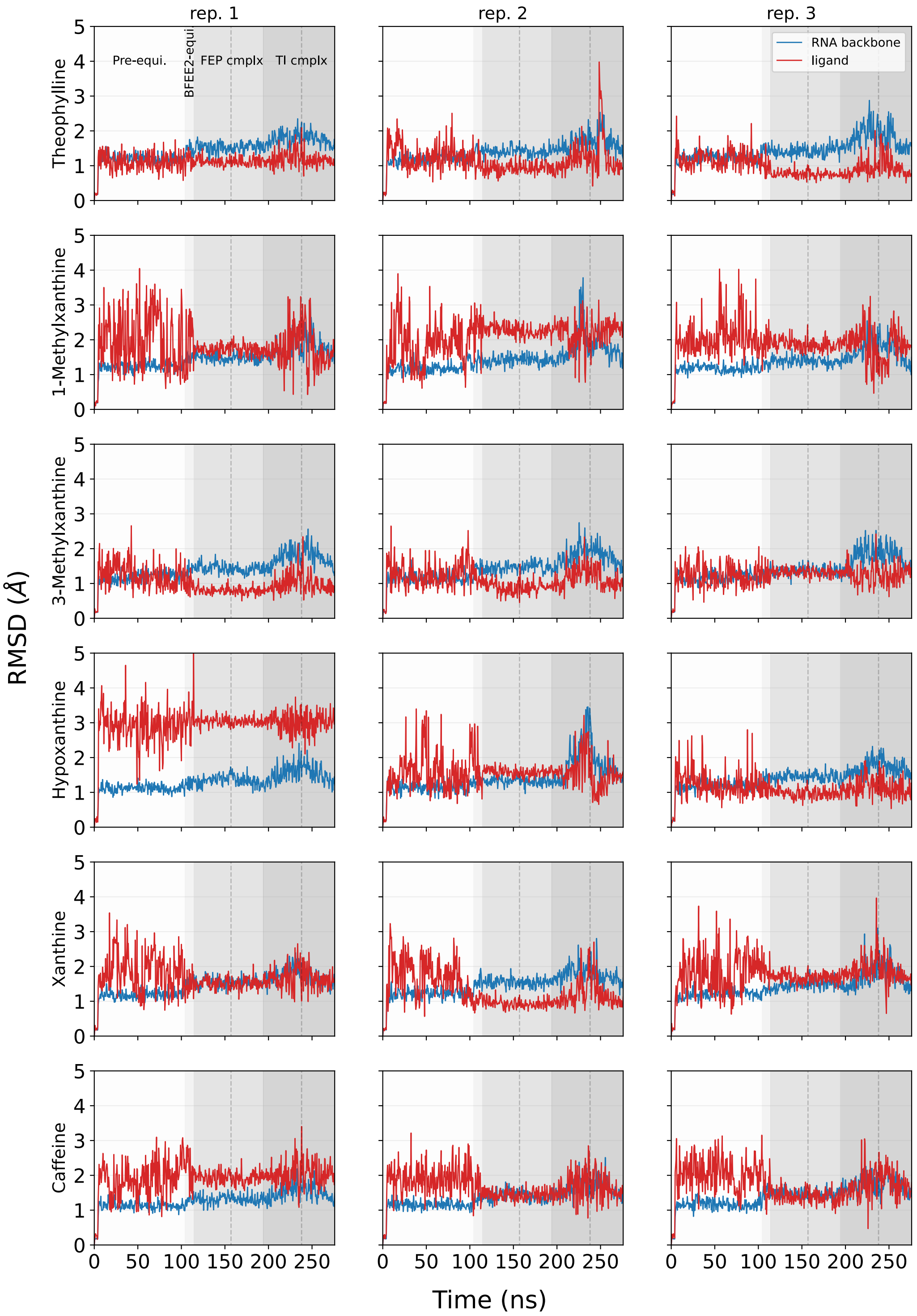


**Figure S15. Stability analysis of the RNA and the bound small molecule for Method 4.** Similar to Figure S12 but for systems with 55 mM NaCl and 0 $Mg^{2+}$ and RNA backbone restraints.

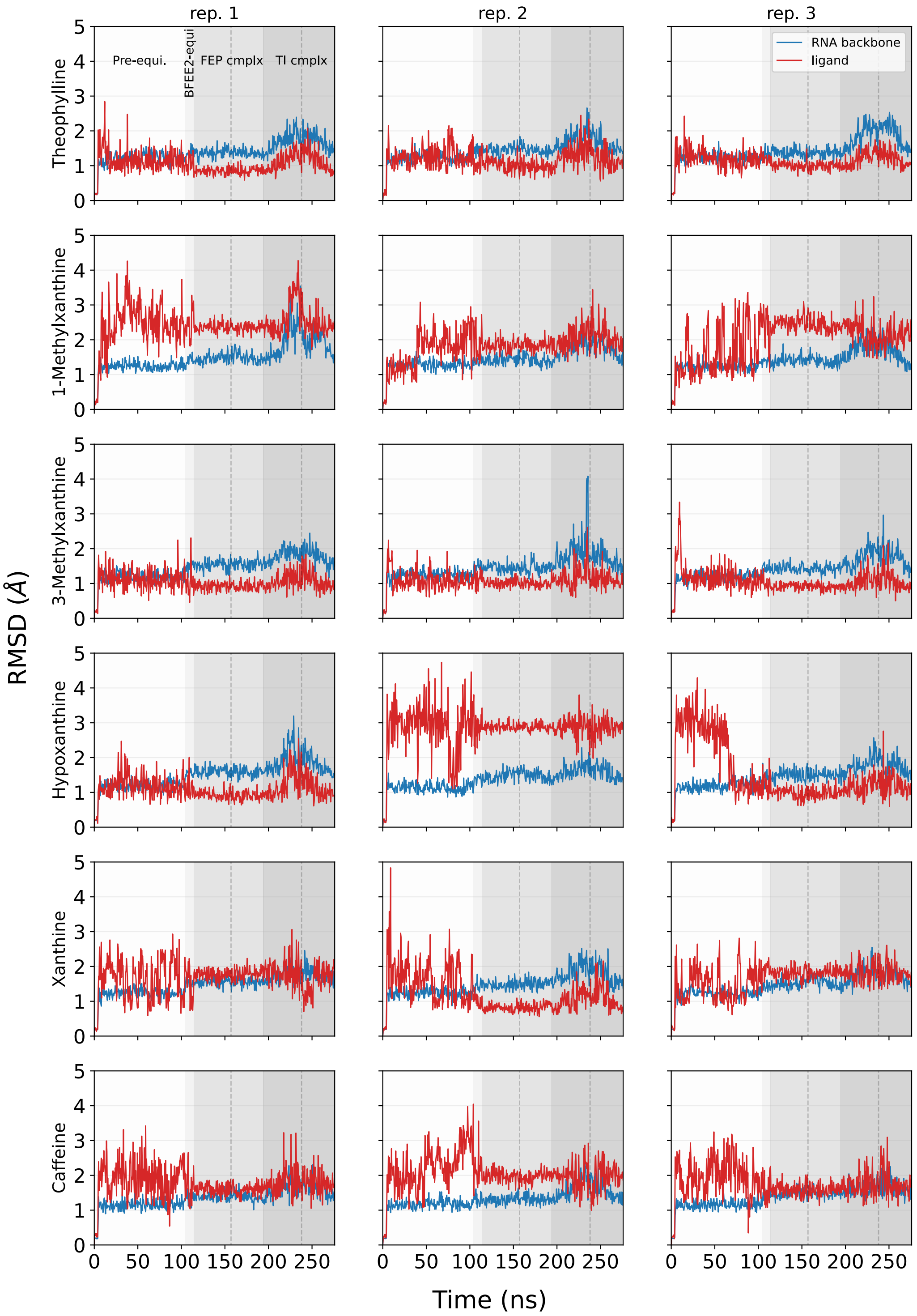


**Figure S16. Stability analysis of the RNA and the bound small molecule for Method 5.** Similar to Figure S12 but for systems with 55 mM NaCl and 2 $Mg^{2+}$ and RNA backbone restraints.

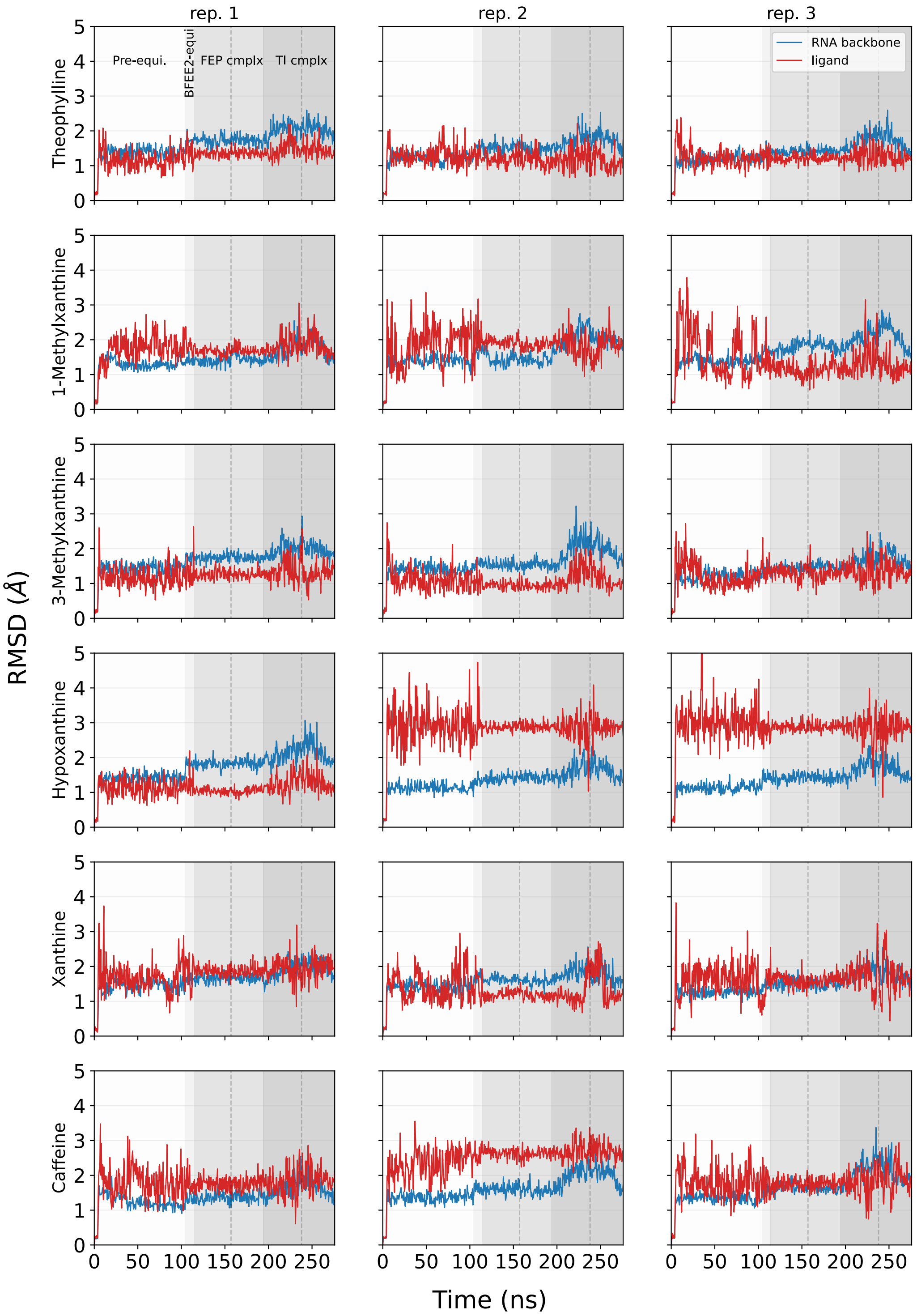


**Figure S17. Stability analysis of the RNA and the bound small molecule for Method 6.** Similar to Figure S12 but for systems with 55 mM NaCl and 3 $Mg^{2+}$ and RNA backbone restraints.

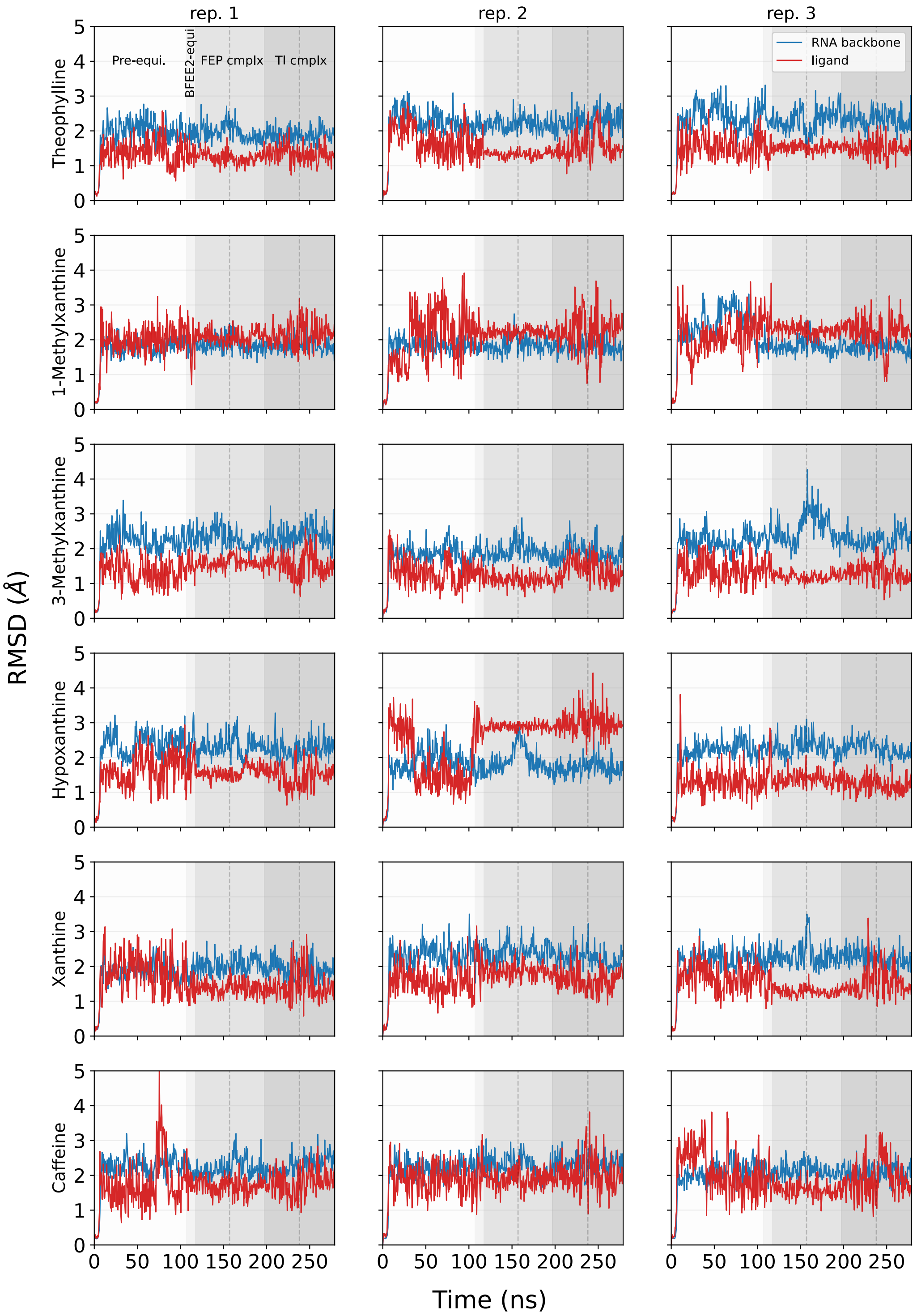


**Figure S18. Stability analysis of the RNA and the bound small molecule for Method 9.** Similar to Figure S12 but for systems with 55 mM NaCl and 3 $Mg^{2+}$ and OpenFF forcefield describing the small molecules' interactions.

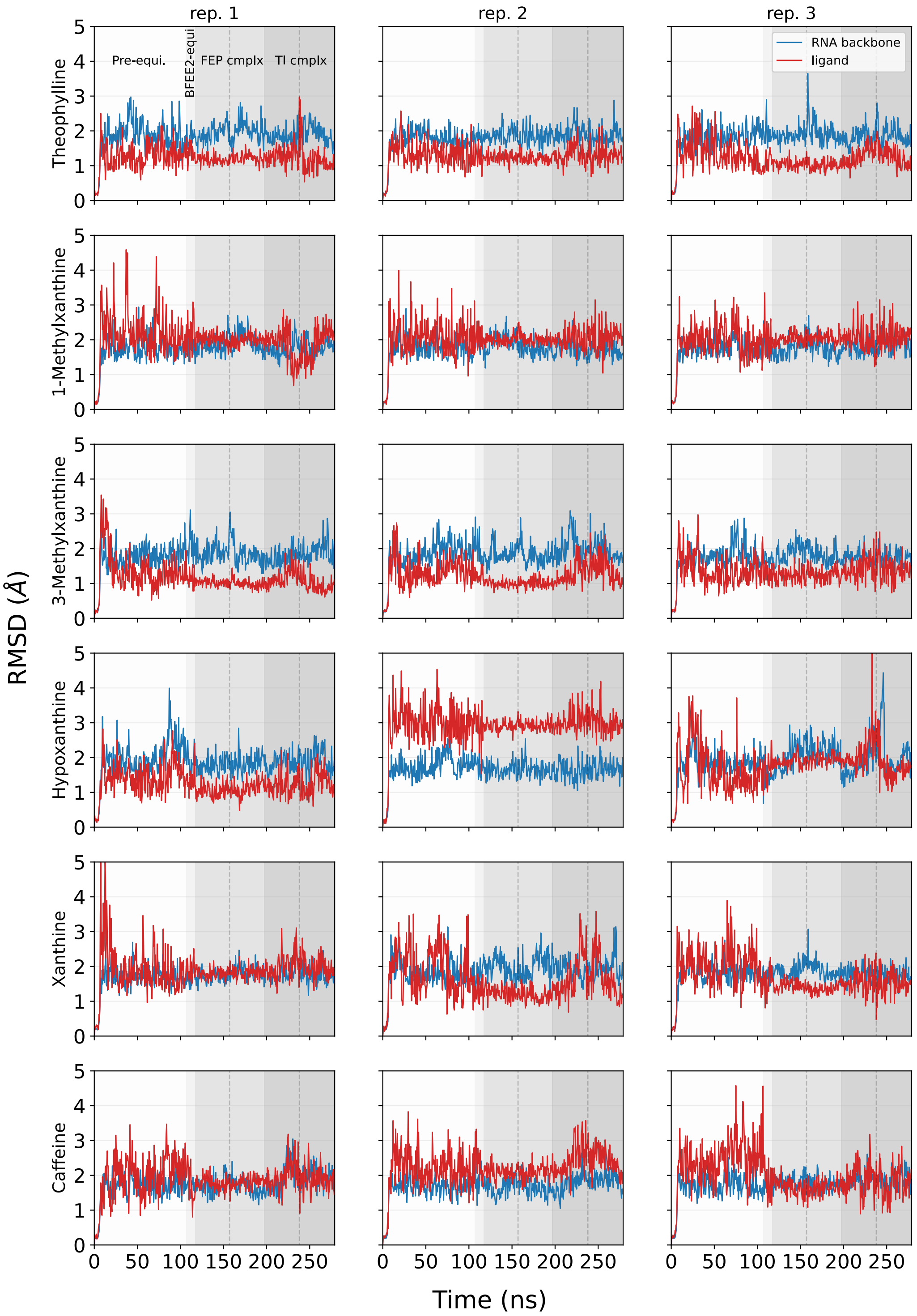


**Figure S19. Stability analysis of the RNA and the bound small molecule for Method 7.** Similar to Figure S12 but for systems with 55 mM NaCl and 3 $Mg^{2+}$ and OPC water model.

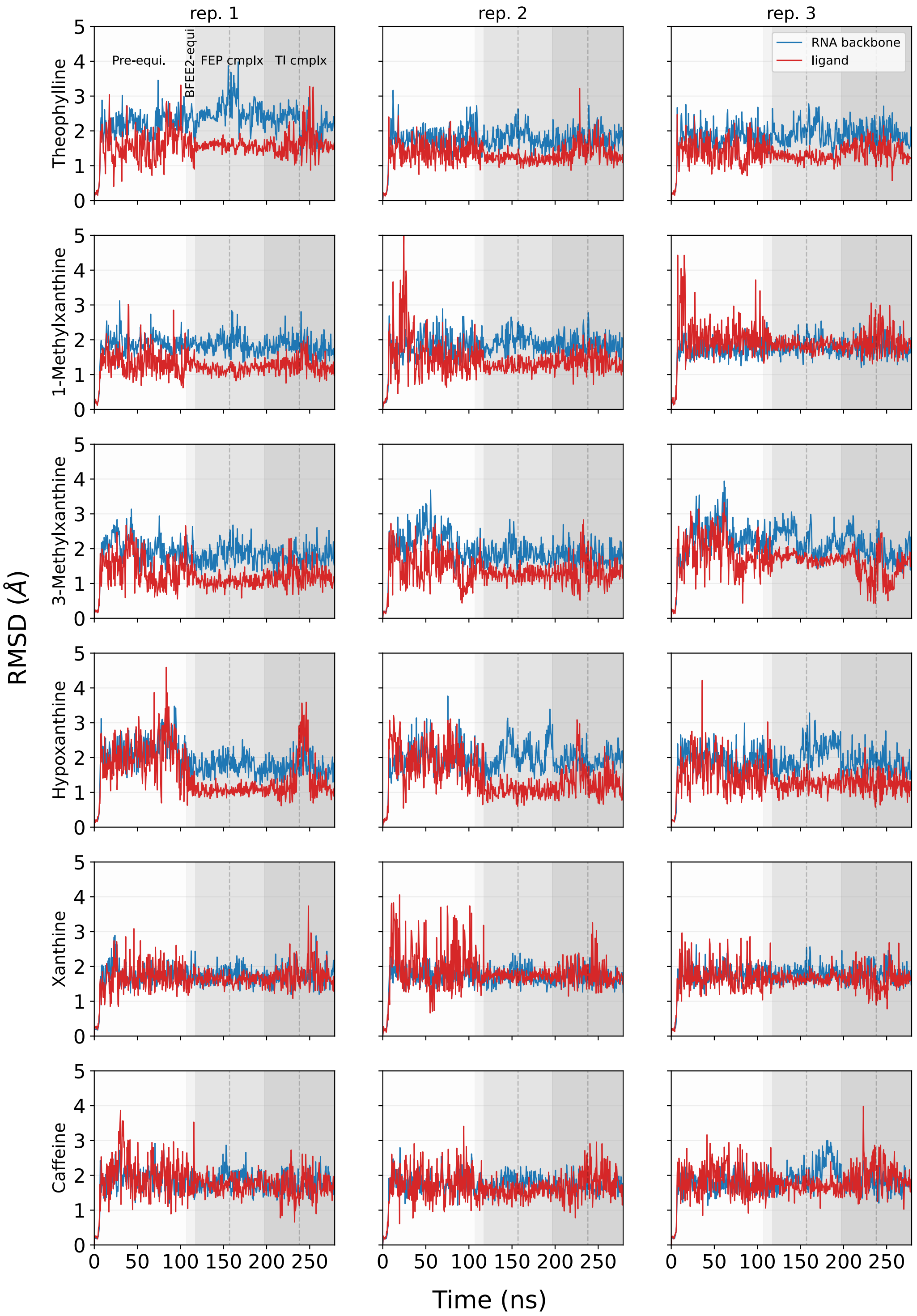


**Figure S20. Stability analysis of the RNA and the bound small molecule for Method 10.** Similar to Figure S12 but for systems with 55 mM KCl and 2 $Mg^{2+}$.

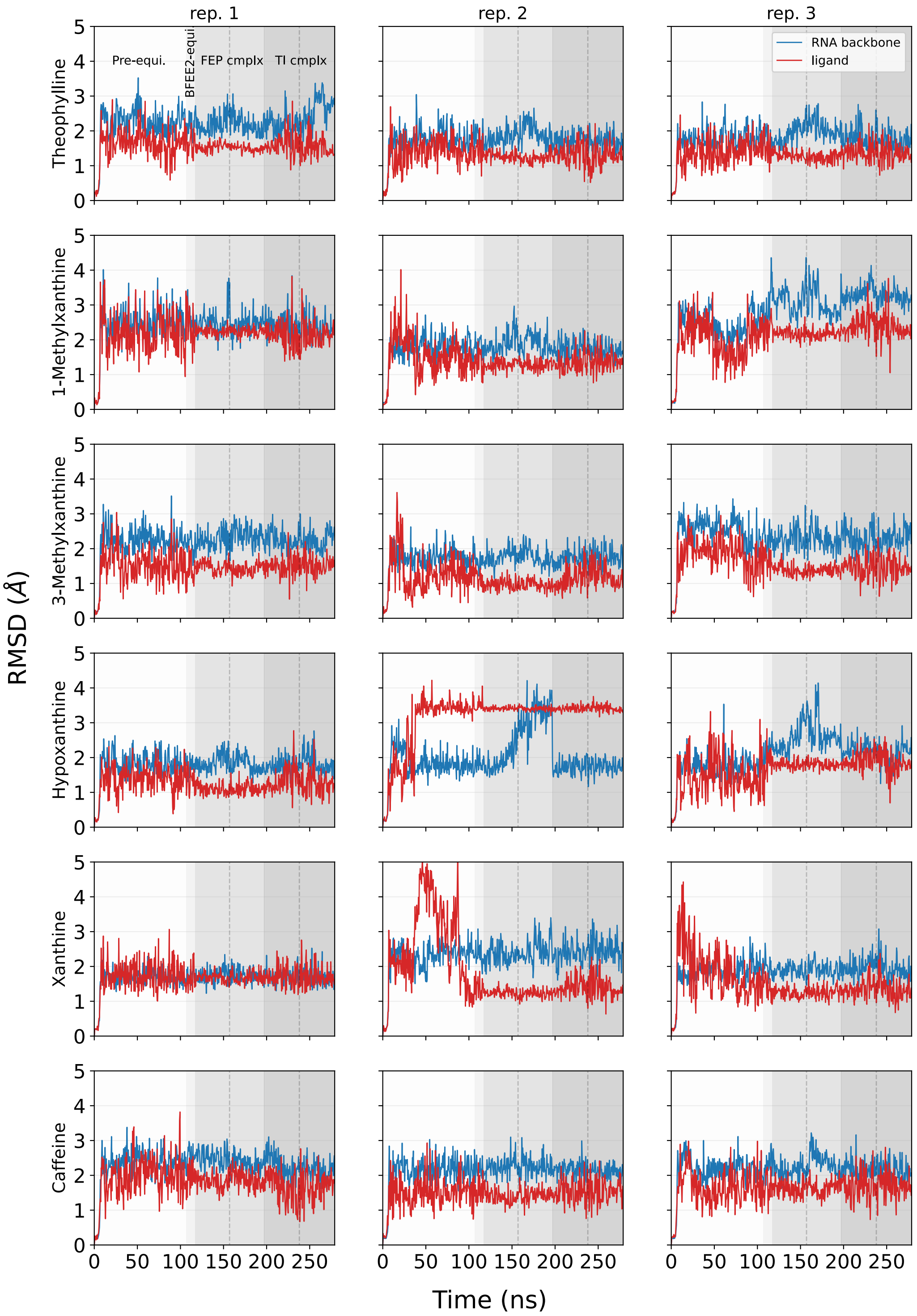


**Figure S21. Stability analysis of the RNA and the bound small molecule for Method 11.** Similar to Figure S12 but for systems with 55 mM KCl and 3 $Mg^{2+}$.

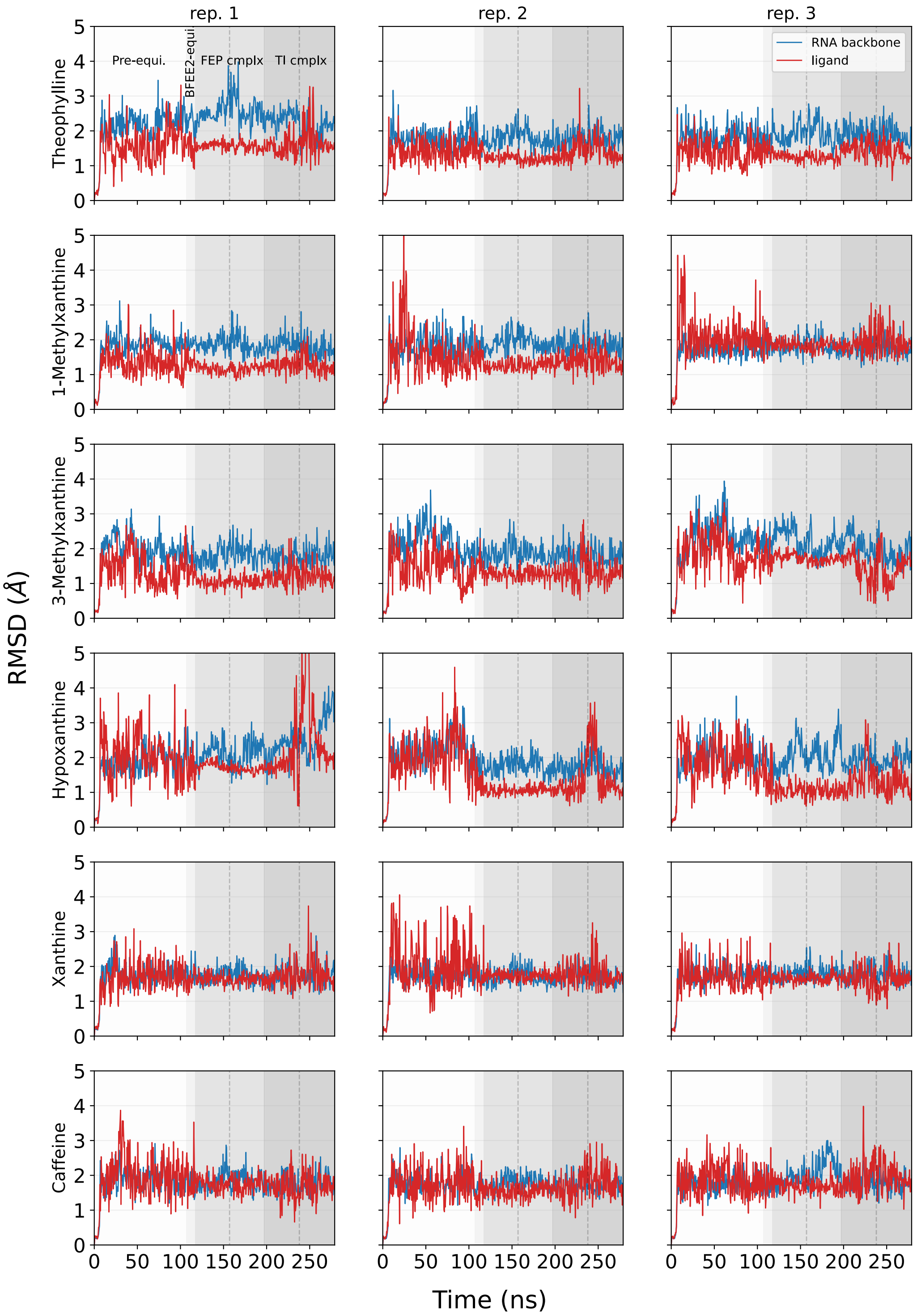


**Figure S22. Stability analysis of the RNA and the bound small molecule for Method 12.** Similar to Figure S12 but for systems with 150 mM KCl and 3 $Mg^{2+}$.

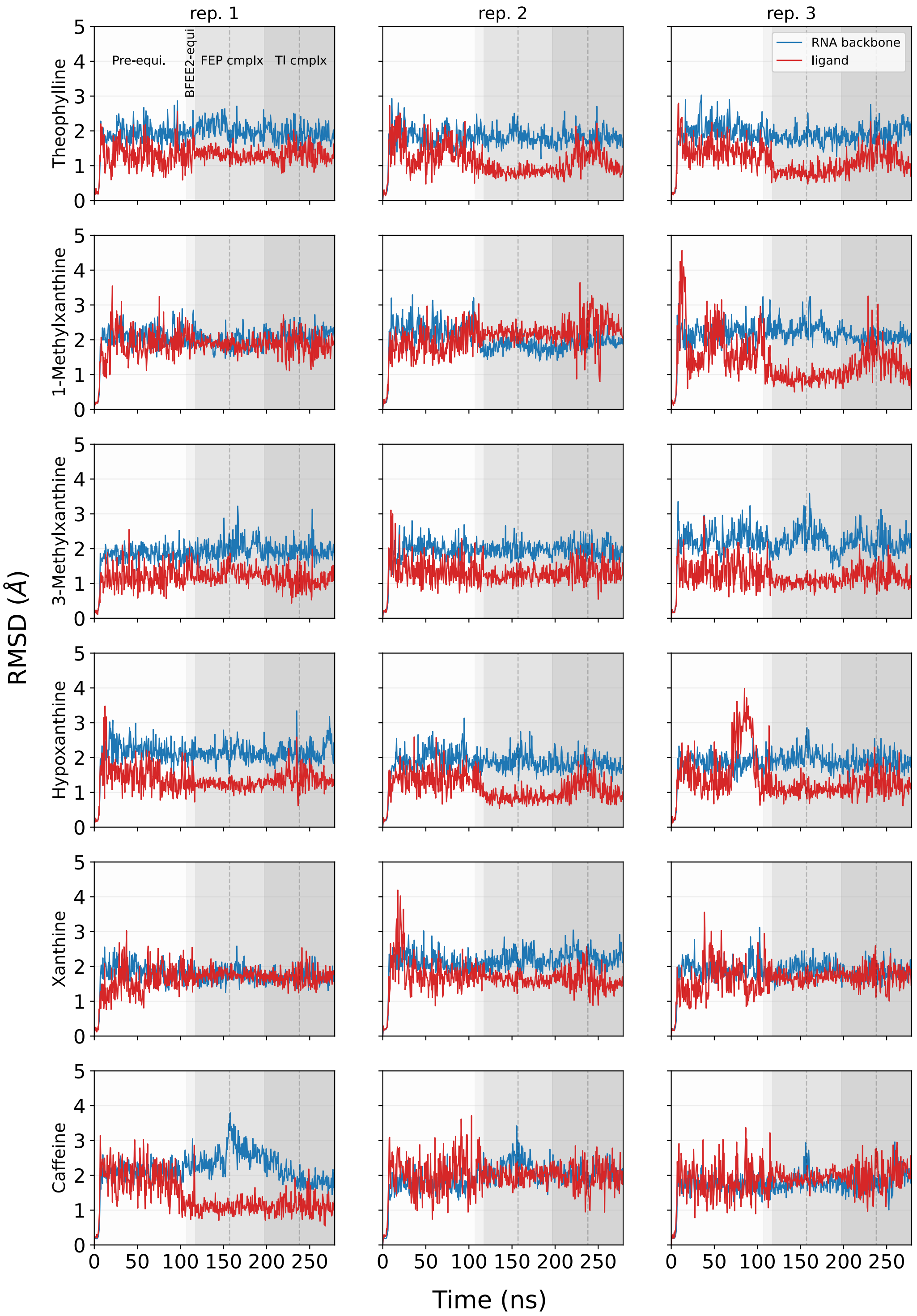


**Figure S23. Stability analysis of the RNA and the bound small molecule for Method 13.** Similar to Figure S12 but for the neutralized systems with 3 $Mg^{2+}$.

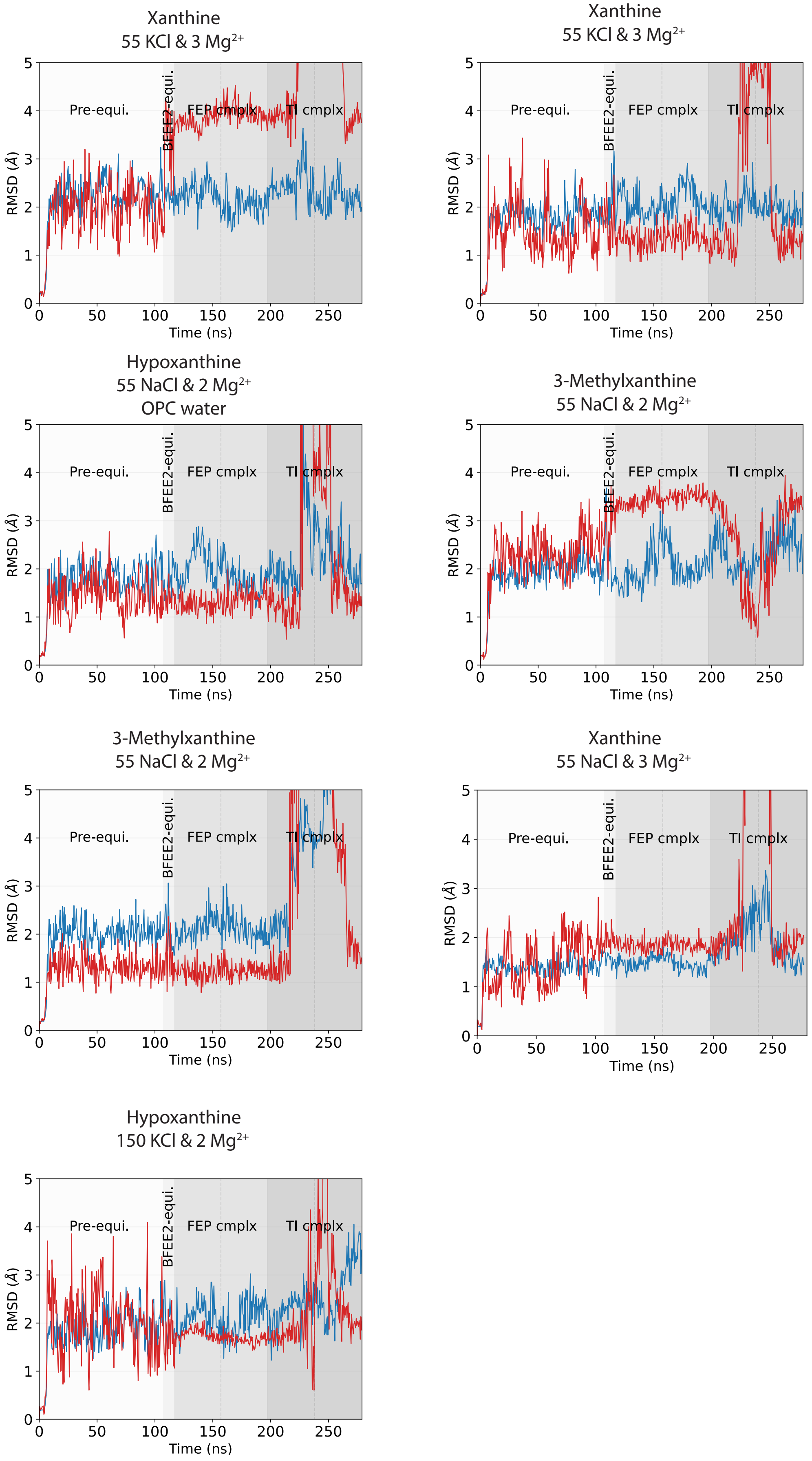


**Figure S24. Stability analysis of the RNA and the bound small molecule in the failed replicas.** Similar to Figure S12 but for the replicas which failed based on the criteria described in Section 2.6 "Rejection protocol for replicate quality control". Red line indicates ligand RMSD and blue line indicates RNA backbone RMSD.
